# Alveolar epithelial progenitor cells drive lung regeneration via dynamic changes in chromatin topology modulated by lineage-specific Nkx2-1 activity

**DOI:** 10.1101/2022.08.30.505919

**Authors:** Andrea Toth, Paranthaman Kannan, John Snowball, J. Matthew Kofron, Joseph A. Wayman, James P. Bridges, Emily R. Miraldi, Daniel Swarr, William J. Zacharias

## Abstract

Lung epithelial regeneration after acute injury requires coordination of extensive cellular and molecular processes controlling proliferation and differentiation of specialized alveolar cells to pattern the morphologically complex alveolar gas exchange surface. During regeneration, specialized Wnt-responsive alveolar epithelial progenitor (AEP) cells, a subset of alveolar type 2 (AT2) cells, proliferate and transition to alveolar type 1 (AT1) cells, though the precise molecular and epigenetic determinants of these processes remain unclear. Here, we report a refined primary murine alveolar organoid assay which recapitulates critical aspects of *in vivo* regeneration, providing a tractable model to dissect these regenerative processes. Clonal expansion of single AEPs generate complex alveolar organoids with extensive structural maturation and organization. These organoids contain properly patterned AT1 and AT2 cells surrounding numerous alveolar-like cavities with minimal structural contribution from mesenchymal cells, implying extensive cell autonomous regenerative function encoded in adult AEPs. Leveraging a time series of paired scRNAseq and scATACseq, we identified the AEP state at single cell resolution and described two distinct AEP to AT1 intermediate states: a widely reported Krt8^+^ transitional state defined by cell stress markers and a second state defined by differential activation of signaling pathways mediating AT1 cell differentiation. Transcriptional regulatory network (TRN) analysis demonstrated that these AT1 transition states were driven by distinct regulatory networks controlled in part by differential activity of Nkx2-1. Genetic ablation of Nkx2-1 in AEP-derived organoids was sufficient to cause transition to a proliferative stressed Krt8^+^ state characterized by disorganized, uncontrolled growth. Finally, AEP-specific deletion of Nkx2-1 in adult mice led to rapid loss of AEP state, clonal expansion, and disorganization of alveolar structure, implying a continuous requirement for Nkx2-1 in maintenance and function of adult lung progenitors. Together, these data provide new insight into cellular hierarchies in lung regeneration and implicate dynamic epigenetic maintenance via lineage transcription factors as central to control of facultative progenitor activity in AEPs.

## Introduction

The pulmonary alveolar gas exchange surface is frequently challenged by pathogens, environmental toxins, and inhaled irritants. These challenges cause chronic, recurrent stress and/or injury to the epithelial, endothelial, and mesenchymal lineages that constitute the alveolus^1^. Homeostatic turnover and regenerative capacity in the alveolus must be sufficient to maintain adequate oxygenation and ventilation throughout life^2,3^. Therefore, extensive recent attention has focused on epithelial progenitor capacity and plasticity in repair of the alveolar epithelium^1,4^ – a topic whose importance has been further emphasized by the COVID-19 pandemic^5,6^. Given the dearth of therapies to promote alveolar epithelial regeneration, models to define epithelial cell relationships and identify pro-regenerative pathways are an area of high priority research interest.

Organoids provide one promising avenue for modeling regenerative biology using adult cells. Multiple lung organoid approaches have been reported in recent years, derived both from primary lung epithelium^7^ and induced-pluripotent stem cells (iPSC)^8-10^. Several themes emerge from these reports. First, the term “lung organoid” encompasses a broad, heterogenous set of cultures with different compositions and morphologies. Second, advancements in purity of epithelial components and/or removal of mesenchymal supportive cells have been reported^11,12^, generally at the expense of complexity. Co-culture systems are characteristically higher in cellular heterogeneity, which is an advantage in replicating the complex cellular composition of the alveolus, but the reproducibility of these co-cultures has been challenged^11^. Third, while iPSC-derived alveolar cells have advanced understanding of human alveolar type 2 (AT2) differentiation and biology^9,13^, it is difficult to model complex adult lung epithelial phenotypes and pathologies using human iPSC cultures due to differences between immature and mature lung epithelium. Finally, lung regeneration involves complex *in vivo* morphogenesis occurring in tandem with cellular differentiation^1^. A major barrier to building an “alveolus in a dish” is the lack of morphological similarity between *in vitro* and *in vivo* models. These challenges have limited the utility of organoid cultures as a method for studying alveolar regeneration *in vitro*.

To address these challenges, we refined and standardized the conditions and inputs of co-culture of murine AT2 cells and alveolar fibroblasts^3,14-16^. Recent data demonstrate that Wnt-responsive AT2 cells, also called alveolar epithelial progenitors (AEPs), harbor extensive progenitor capacity^3,17,18^. Following injury, AEPs expand rapidly, differentiate into new AT1 and AT2 cells, and repair regions of alveolar injury following epithelial loss or infectious stress. Herein, we describe the high dimensional characterization of these organoids across multiple timepoints. We find that AEP-derived organoids, or AEP-O, develop via clonal expansion of single progenitor cells and undergo progressive cellular differentiation and spontaneous cavity formation *in vitro*. Using multistage single cell transcriptomics and epigenomics, we defined the organoid cellular milieu and identified separable AEP, AT2, AT1, and transitional states^19-22^ in organoids. We validated these states based on published *in vivo* scRNAseq and derived cellular trajectories and lineage relationships from a known progenitor root state. Comparative transcriptional regulatory network (TRN) analysis^23,24^ along these trajectories identified several known and novel regulators of alveolar epithelial biology and highlighted a novel role for the lineage transcription factor Nkx2-1^25-27^ in progenitor and transitional lineages of the adult alveolus. Genetic ablation of Nkx2-1 in AEPs *in vitro* and *in vivo* caused irreversible acquisition of a stressed transitional state, with uncontrolled proliferative growth and disruption of organoid morphology *in vitro* and alveolar structure *in vivo*. These findings highlight the utility of the AEP-O assay as a high-fidelity model of *in vivo* alveolar biology and implicate Nkx2-1 as a central regulator of alveolar epithelial progenitors.

## Results

### Alveolar epithelial progenitor-derived lung organoids recapitulate key cellular and morphological aspects of alveolar regeneration

We generated AEP-O by combining FACS-sorted AEPs derived from Axin2^CreERT2-tdT^ mice at a 1:10 ratio with lung mesenchymal cells isolated by selective adhesion in a 1:1 mix of Matrigel (Corning) and small airway growth media (SAGM, Lonza). To evaluate the internal growth and maturation of these organoids, we adapted methods from iPSC culture for whole mount immunohistochemistry of organoids^28^ and performed a time series evaluation using high content confocal imaging paired with longitudinal single cell sequencing (Figure 1A).

**Figure 1.**
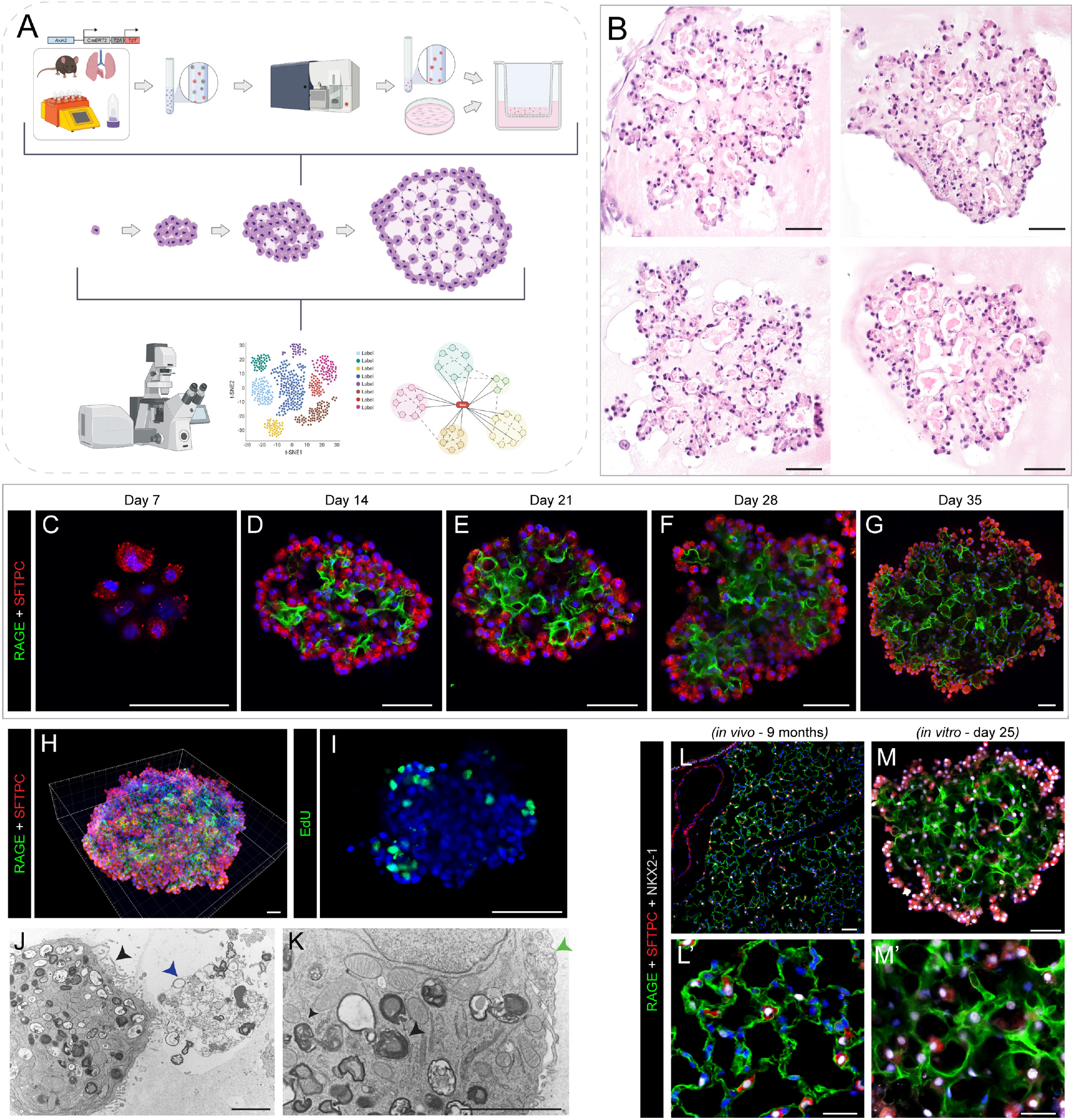
AEP-derived alveolar organoids clonally expand and pattern complex, polarized alveolar-like cavities. (A) Schematic of experimental design and overview. Live/CD31^-^/CD45^-^/CD326^+^(EpCAM^+^)/TdTomato^+^(Axin2^+^) cells (AEPs) were mixed with mouse lung fibroblasts from P28 mice and cultured for up to 35 days, followed by analysis via high content imaging. (B) H&E of 5 μm sections of FFPE day 35 Axin2^+^ organoids, showing cellular morphologies typical of both AT1 and AT2 cells. (C-G) Whole-mount immunofluorescence time course of Axin2^+^ organoids showing expansion of SFTPC^+^ AT2 cells (red), increased differentiation into RAGE^+^ AT1 cells (green) and increased structural complexity. (H) Imaris 3D reconstruction of day 35 Axin2^+^ organoid (z-depth = 174.13 μm) showing cellular arrangement/organization within mature organoids. (I) Click-iT EdU (green) whole-mount day 25 Axin2^+^ organoids, with proliferating cells primarily on outer edges or ‘buds’ growing outward from the organoid. (J-K) Electron microscopy of day 28 organoids. (J) Image of properly polarized AT2 cell with apical microvilli (black arrowhead) secreting surfactant (blue arrowhead) into a lumen. (K) Image of AT2 cell with lamellar bodies (black arrowhead) adjacent to an AT1 cell (green arrowhead, right). (L-M) Comparison of *in vivo* mouse lung (9-month C57BL/6J mouse) and *in vitro* day 25 Axin2^+^ organoids. [*Scale bars = 50 μm, except for electron microscopy (J, K) scale bars = 2*.*5 μm*]. (*RAGE = Receptor for Advanced Glycation End-products [AT1 cell marker]; SFTPC = Surfactant Protein C [AT2 cell marker]; EdU = 5-ethynyl-2’-deoxyuridine; FFPE = formalin-fixed, paraffin-embedded*)

AEP-O grow clonally from a single AEP in definable stages. First, AEPs expand into small clusters of SFTPC^+^ cells during the first week of culture (Figure 1C). By day 14, differentiation of RAGE^+^ AT1 cells was observed within the central portion of the organoids (Figure 1D), consistent with prior reports^3,14-16,19,21,29^. During the third week of culture, these developing AT1 cells elongate and polarize (Figure 1E), and by d28 in culture cavities were present within the organoids (Figure 1B, F). These cavities matured into a network of alveolar-like structures during the 4^th^ and 5^th^ weeks of differentiation (Figure 1F-G). AT1 cells intermixed with AT2 cells within the central portion of the organoid (Figure 1H, M), with minimal apoptosis detectable by TUNEL staining in AEP-O during cavity formation (Figure S1A-D). Continued proliferation was evident by EdU incorporation at the periphery of mature organoids (Figure 1I). Electron microscopy demonstrated that the epithelial lining of these cavities includes AT2 cells containing lamellar bodies with the apical surface directed towards the internal lumen and evidence of active surfactant secretion adjacent to elongated AT1 cells (Figure 1J-K). These features of the mature cavities within AEP-O bear a striking similarity to the epithelial structure of mature murine alveoli (Figure 1L-M), suggesting an unprecedented degree of morphological maturation in AEP-O compared to other reported lung organoids.

### AEP-O maturation is driven by mesenchymal paracrine signaling without direct mechanical contribution of mesenchymal cells

To better characterize the progressive cellular maturation occurring during paired cell differentiation and cavity formation in AEP-O, we performed single cell RNA sequencing at 14, 21, and 28 days after culture initiation. We identified epithelial and mesenchymal populations, as well as an unexpected immune fraction (Figure 2A-C)^30^. Epithelial contribution increases from d14 to d28, with progressive maturation and increase in relative proportion of alveolar type 1 cells (Figure 2B, Figure S2A-D). To evaluate the determinants of epithelial maturation in AEP-O, we examined the supportive cells comprising the signaling niche in these organoids.

**Figure 2.**
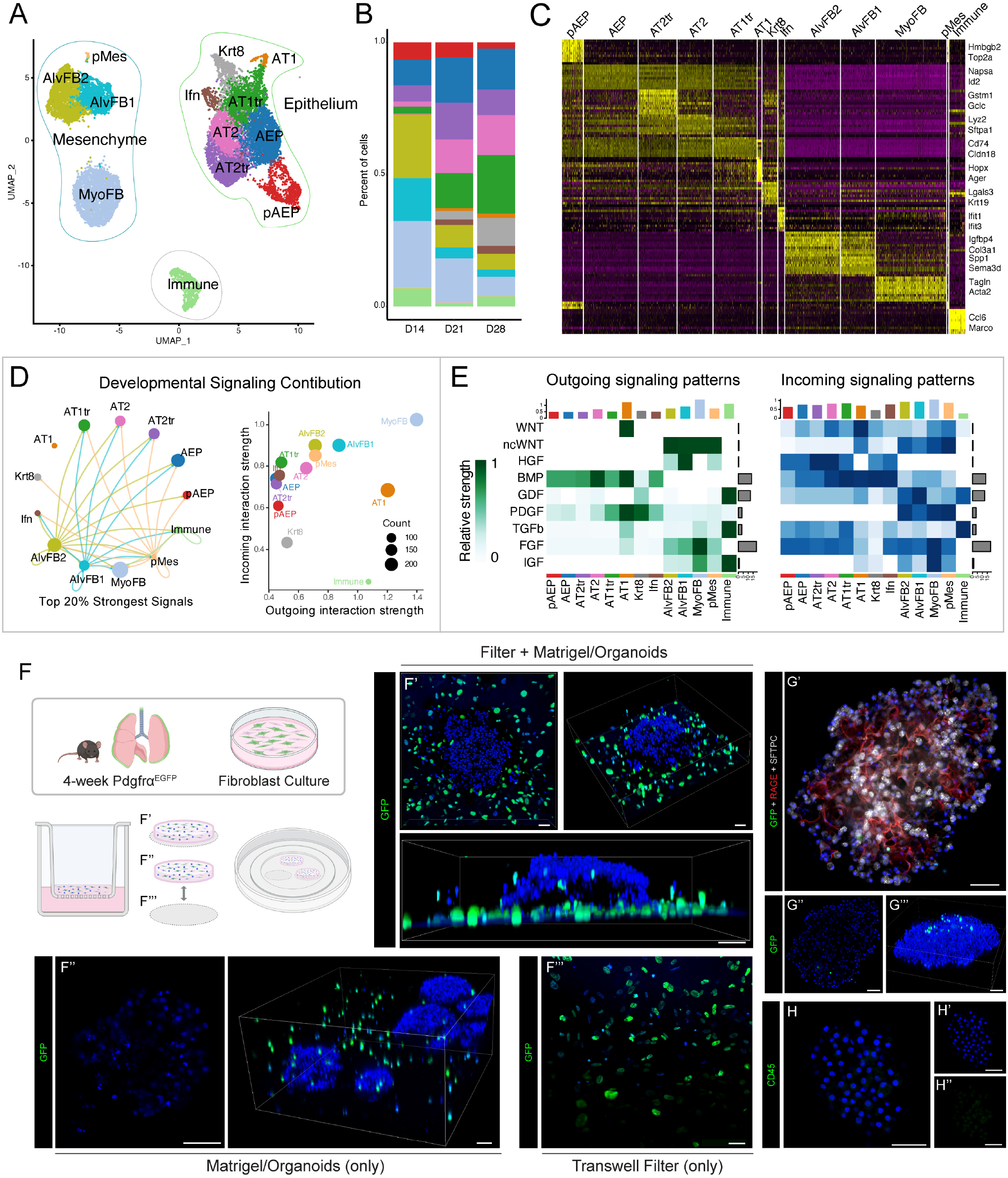
Single cell composition and epithelial-mesenchymal interactions in alveolar organoids over time course of differentiation. A) UMAP of all cell populations combining d14, d21 and d28 AEP-derived organoid scRNAseq datasets. B) Cell population proportions at each time point with increasing proportion of epithelial cells. C) Heat map showing expression of top 10 most differentially expressed genes in each population D-E) Ligand-receptor analysis of organoid culture demonstrating extensive mesenchymal-epithelial communication in organoids. F) Schematic of experimental set-up of live imaging and 3D reconstruction of live day 20 organoids generated using PDGFRα^EGFP^ fibroblasts stained with Hoechst, with data shown in F-G. F) 3D reconstruction of confocal z-stacks of whole wells including Transwell filter, showing the majority of GFP^+^ fibroblasts are growing on the filter; F’-F’’’) Whole-mount immunofluorescence showing lack of PDGFRα^+^ cells within day 20 organoids, with scattered cells found throughout the surrounding Matrigel. G) Whole mount IHC showing few GFP^+^ fibroblasts inside of organoids. H) CD45 staining of organoids; see also Figure S3I. [*Scale bars = 50 μm*]; *(PDGFRα = platelet-derived growth factor receptor alpha; GFP = Green Fluorescent Protein; RAGE = Receptor for Advanced Glycation End-products [AT1 cell marker]; SFTPC = Surfactant Protein C [AT2 cell marker]); AEP = alveolar epithelial progenitor, pAEP = proliferative AEP, AT2tr = AT2 transitional cell, AT2 = alveolar type 2 cell, AT1tr = AT1 transitional cell, AT1 = alveolar type 1 cell, Krt8 = Krt8/DATP/PATS-like transitional cell, Ifn = Interferon responsive alveolar cell, pMes = proliferative mesenchymal cell, AlvFB1 = alveolar fibroblast type1, AlvFB2 = alveolar fibroblast type 2, SM = smooth-muscle like mesenchyme*.

Within the mesenchyme, two major PDGFRα^+^ populations^14,15,31-33^ were identified, corresponding to murine alveolar fibroblasts and myofibroblasts in LungMAP datasets^31^. To localize these mesenchymal cells in complex culture, PDGFRα^EGFP^ lung fibroblasts^34^ were obtained via selective adhesion (Figure S3A-B). Use of these PDGFRα^EGFP^ fibroblasts did not affect organoid growth (Figure S3C-D). PDGFRα^EGFP^ cells localize predominantly in two locations, with the minority of cells surrounding the epithelial organoids suspended in Matrigel, and the majority growing on the Transwell filter in a monolayer (Figure 2F-H, S3E-F). Few PDGFRα^EGFP^ cells were detected within organoids (Figure 2F-G) and no clear deposition of fibrillar collagen (types I and II) was seen within organoids (Figure S3H), suggesting that the morphological maturation and complex structural organization of AEP-O did not require direct mesenchymal cell localization within the organoid itself. CD45 staining showed that immune cells were scattered throughout the fibroblast stocks and within the Matrigel, but never found in large clusters or within organoids (Figure S3I); we concluded that these represented a minor contaminant of the supportive fibroblasts. Therefore, we hypothesized that the mesenchymal cells were predominantly involved in providing a signaling niche in AEP-O.

Ligand-receptor analysis^35^ suggested extensive signaling between epithelial and mesenchymal cells within AEP-O (Figure 2D-E). Major signal producers in AEP-O included AT1 cells, matrix fibroblasts and myofibroblasts. AT1 cells expressed WNT ligands with predicted receptivity in both WNT-responsive AT2 cells and multiple mesenchymal populations. AT1 also produced PDGF ligands predicted to signal to the PDGFRα^+^ mesenchyme. Mesenchymal cells expressed HGF, non-canonical WNT, and FGF ligands, consistent with published data describing roles of these pathways in alveolar regeneration^15,26,36-41^. Together, these data suggested that the AEP-O signaling milieu recapitulates key aspects of the *in vivo* regenerative niche, and that the mesenchymal cells provided a supportive paracrine signaling niche required for alveolar cavity formation.

To directly evaluate the requirement for mesenchymal signaling support, we cultured of PDGFRα^EGFP^ cells on the basolateral side of the Transwell filter (Figure S3G). Absence of fibroblasts in the Matrigel plug led to complete loss of organoid formation (Figure S3G), confirming a requirement for paracrine mesenchymal signaling in the establishment and maturation of AEP-O.

### scRNAseq defines separable epithelial maturation trajectories of AEPs toward AT1 and AT2 cells within alveolar organoids

Integrated analysis of data from d14, 21, and 28 identified eight separable epithelial cell states via graph-based clustering in Seurat^30^ (Figure 2A-C, Figure S2A-D). Consistent with our whole mount IHC results, both AT1 and AT2 cells were identifiable within organoids. We noted a clear AEP state defined by expression of the AEP-enriched markers *Id2, Ctnnb1, Lrp5, Lrp2, Napsa, Bex2, Hdc*, and *Fgfr2*^*3,17*^. Cells bearing the AEP signature were also found in a second state partially defined by high level expression of cell cycle genes. Given that the starting epithelial fraction was sorted AEPs, we called these cells AEPs and proliferative AEPs (pAEPs), respectively (Figure 2A). The proliferative state is readily detectable in day 14 cultures and decreases by day 28 (Figure 2BA-C, S2A-D); EdU staining of organoids confirmed a proliferative Sftpc^+^ positive population at day 25 (Figure 1I).

To dissect pathways of differentiation from the AEP state, we compared lineage predictions generated by both trajectory analysis^42^ and RNA velocity^43^ (Figure 3A-B). Differentiation of AEPs toward AT2 cells was detected through an AT2 transitional state (AT2tr). The AT2tr state is defined by high level expression of glutathione pathway genes and shift towards lipid metabolism, while the AT2 state expressed high levels of mature AT2 markers including *Sftpa1* and *Lys2* (Figure 3C). Other AT2 markers, including *Sftpc* and *Sftpb*, were expressed broadly within multiple differentiating AEP states (Figure 2C, 3F).

**Figure 3.**
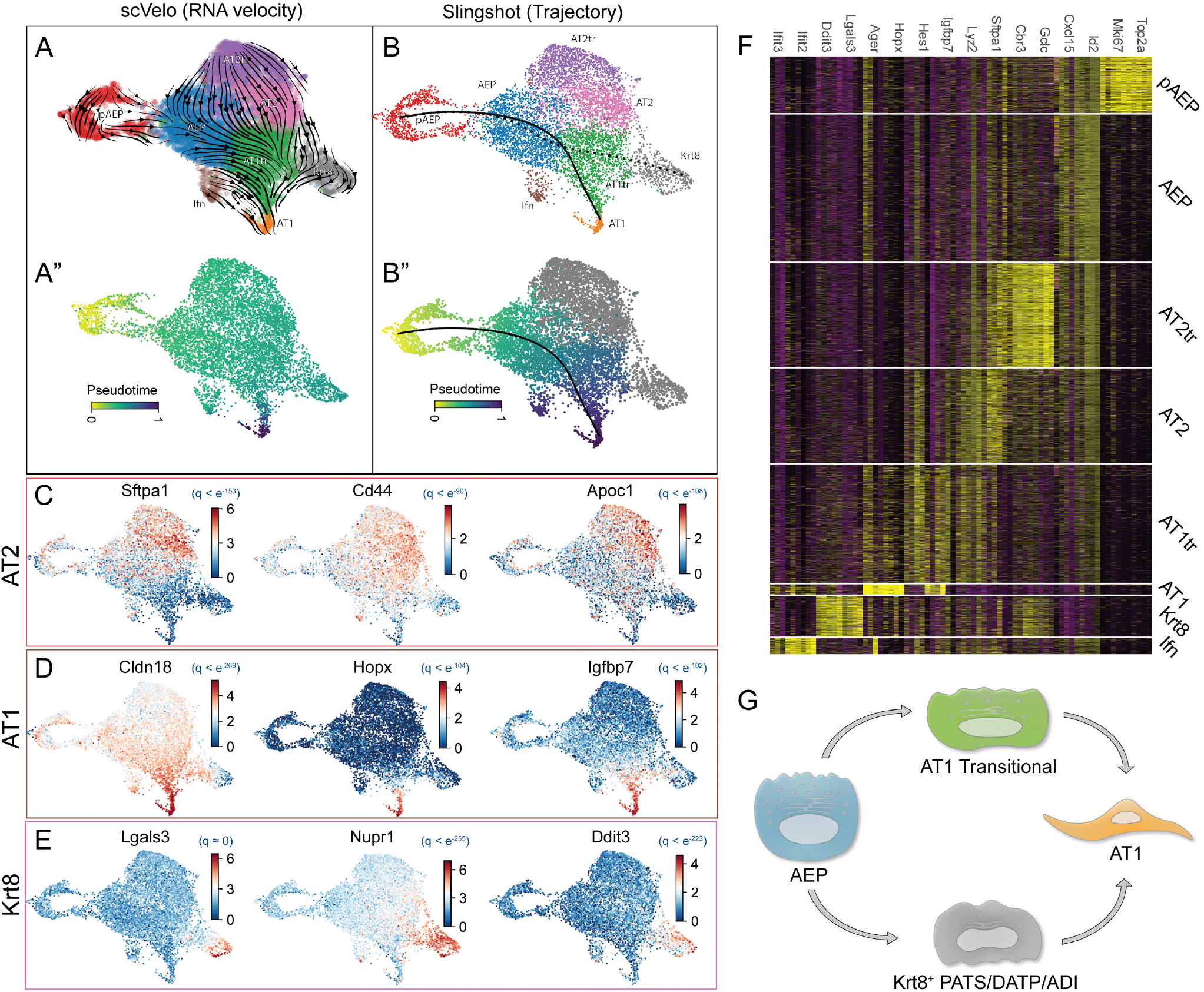
AEP-derived organoids elucidate dynamics of alveolar epithelial differentiation. A) scVelo RNA velocity UMAP showing differentiation dynamics (A) and pseudotime inferred from RNA velocity (A”) in AEP organoids. B) Slingshot trajectory analysis and pseudotime inference of AEP organoids demonstrates similar lineage relationships to RNA velocity. C-E) Lineage drivers defined by CellRank for differentiation of pAEP/AEP to AT2 cells (C), AT1 cells via AT1tr (D), and Krt8 cells (E). F) Heatmap showing major cell markers differentiating cell states in alveolar epithelium. G) Model of cellular relationships and AT1 differentiation inferred from alveolar organoids.

We next examined AT1 differentiation in AEP-O. Recent attention has focused on differentiation of AT2 progenitor cells to AT1 cells, with multiple reports describing a distinct transitional state (variously called PATS^19^, DATP^21^, ADI^20^, or Krt8^+^ cells^22^) marked by high level expression of cell stress markers including *Krt8, Lgals3, Tp53*, and *Cldn4*; we will call this stressed cell state Krt8^+^ in this manuscript. Both trajectory analysis and RNA velocity suggested two trajectories from AEP to AT1 cells; one through this Krt8^*+*^ transitional state, and another through a novel second, less stressed state we denoted AT1 transition (AT1tr).

RNA markers of the AT1tr state included *Hes1* and *Igfbp7*, and RNA velocity analysis showed decreasing AT2 gene expression and intermediate expression of AT1 markers within this population (Figure 3D-F), supporting the transitional nature of this cell state. Label transfer and integration with published organoid data sets^19,21^ showed both AT1tr and Krt8^+^ cell states were detectable in published data (Figure S4A-C). These findings suggested the model that AEPs could differentiate toward AT1 cells through either the Krt8^+^ state or the AT1tr state (Figure 3G), rather than through an obligate intermediate state. However, given the challenges of cell state predictions and transitions based on solely on RNA transcriptome^44^, we proceeded to evaluate epigenomic state and chromatin topography of the cells comprising AEP-O via scATACseq.

### AEP chromatin state is defined by a progenitor-enriched transcriptional regulatory network

We performed scATAC sequencing at d14, d21, and d28 in AEP-O, and performed unbiased cell clustering of epigenomic states using ArchR^45^. The same number of epithelial cell states were detected by scRNAseq and scATAC using default parameters, supporting the conclusion that clustering parameters were appropriate in both assays (Figure 4A). Concordant with published bulk ATACseq data^3^, AEP and AT2 are distinct at the epigenomic level (Figure 4B); AT1tr and Krt8^+^ also showed differential chromatin accessibility, with distinct regions of open chromatin (Figure 4C). Integrated analysis combining both scRNA and scATACseq identified seperable clusters of regulated genes within AEPs, for AT2 differentiation, and for AT1 differentiation (Figure 4D). scATACseq-based pseudotime inference confirmed the presence of two separate differentiation trajectories from AEPs to AT1 cells, one passing through the Krt8^+^ state and the second passing through the AT1tr state (Figure 4E-F). Comparison of expression and chromatin accessibility using combining gene expression with ATAC suggested overlapping gene sets shared by Krt8^+^ and AT1tr cells, but also substantive differences (Figure 4D), implying independent regulatory inputs to these two states.

**Figure 4.**
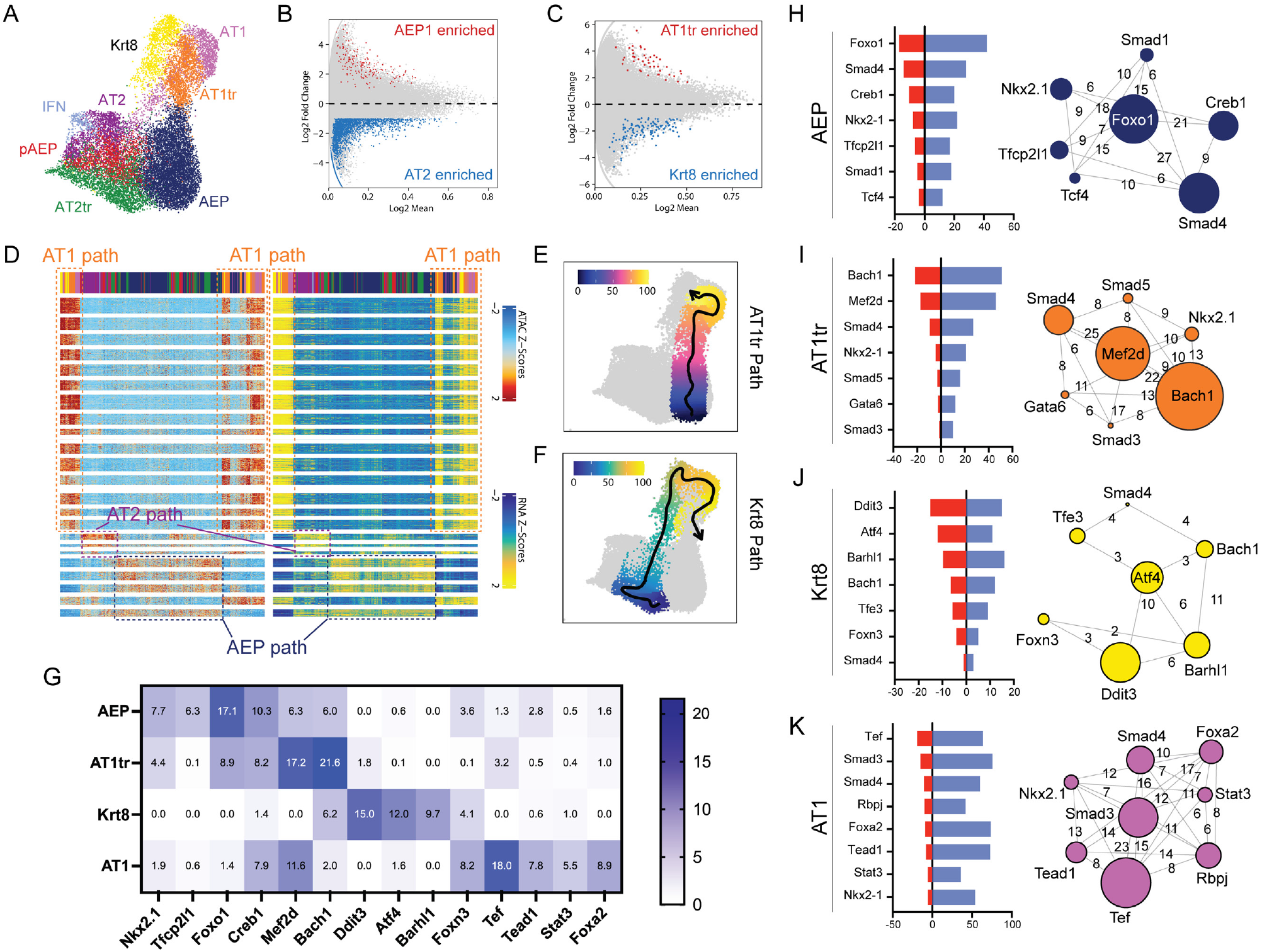
scATACseq analysis of AEP-derived organoid formation. A) UMAP of cellular populations within organoids, named as in RNAseq. B-C) Volcano plots showing differential chromatin accessibility regions between AEP and AT2 cells (B) and AT1tr and Krt8^+^ transitional cells (C). D) Paired heatmap of differentially accessible genomic loci in ATAC (left) and RNA expression of nearest-neighbor gene production (right) showing overview of regulators of AT1 cell differentiation (AT1 path), AT2 cell differentiation (AT2 path), and AEP state (AEP path) derived from integrated analysis. Cell populations shown along top bar, with colors the same as in (A). E-F) Pseudotime prediction of separate AT1 differentiation trajectories from AEPs to AT1 cells through AT1tr path (E) and Krt8^+^ path (F). G) Transcriptional activity score (TAS, negative -log p value of TF enrichment per cell type by Fisher exact test), for TFs per cell type in AT1 differentiation trajectories. H-K) Top transcriptional regulators of AEP (H), AT1tr (I), Krt8^+^ (J), and AT1 (K) cell states. Red bar = TAS, and blue bar = # of regulated genes expressed in given cell type. Network diagram shows core regulator relationship, with circle size indicating TAS, and numbers of bars showing co-regulated gene networks per cell type.

### AEP to AT1 differentiation occurs through multiple separable progenitor states with independent TRNs

To better define the differences between Krt8^+^ and AT1tr states, we turned to transcriptional regulatory network (TRN) inference^23,24^. Transcriptional regulatory network inference models describe interactions between transcription factors and their gene targets within specific cell types, providing predictions relevant to both the driving TFs and the regulated gene sets regulated in a given cell type ^46^. Combination of RNA expression and chromatin accessibility can improve TRN predictions by reducing both false positive and false negative regulatory predictions^47^; application to single cell techniques have extended the power of these approaches to estimate TF regulators of individual cell states^24^. Therefore, we performed TRN inference comparing regulatory networks of various cell states within AEP-O to identify differential TF activity and predict regulators of state transitions (Figure 4G-K). Focusing on the core TFs within each TRN, we found that AEP regulators included *Nkx2-1* and *Tfcp2l1*^*48*^, both enriched in expression in bulk RNAseq from AEPs^3^, as well as TFs modulating Wnt (*Tcf4*) and BMP (*Smad1/4*) activity concordant with known AT2 progenitor signaling response^49-51^ (Figure 4H). Within AT1 cells, we identified *Nkx2-1, Gata6*^*52*^, and *Foxa2*^*53,54*^, all well-known modulators of AT1 gene expression, and signaling response from the AT1-associated Yap/Taz^25,55,56^ (*Tead1*), Notch (*Rbpj*)^57,58^, and Tgfβ^20,59^ (*Smad3/4*) signaling pathways (Figure 4K). These results provided evidence that TRN inference had successfully identified known regulators of the AEP and AT1 cell states, supporting the notion that TRN inference could distinguish factors driving AT1 differentiation.

We therefore focused our attention on differential regulators of the AT1tr and Krt8^+^ states. We calculated transcription factor activity scores based on enrichment in predicted cell type. Both states showed overlap with AT1 enriched TFs, though several of the AT1 TFs were enriched in only one of the transition states. The TRN predictions were otherwise distinct, suggesting differential inputs to the two transitional states. The top regulators in Krt8^+^ cells were *Atf4* and its target *Ddit3*, the gene encoding C/EBP homologous protein (CHOP)^60^ (Figure 4J); together these factors are activated by the multiple inputs of the integrated stress response (ISR)^61,62^, with CHOP implicated in regulation of checkpoints in apoptosis vs cellular differentiation in other systems^60^. The ISR is activated in alveolar epithelium following ventilator-induced lung injury^63^, and ISR activation contributes to lung fibrosis^64^; Atf4 and CHOP activation may underlie Krt8^+^ cell accumulation in fibrosis^19^.

Conversely, the AT1tr TRN is characterized by homeostatic lung epithelial differentiation factors. *Gata6*, a known direct activator of the AT1 differentiation program^52^, is predicted to co-regulate a number of genes in cooperation with Smad4 and Nkx2-1. Chromatin topology suggested that AT1tr cells balance BMP- and Tgfβ-associated SMAD activity (Figure 4I, S5), consistent with previous reports demonstrating a transition from BMP to Tgfβ signaling in AT1 differentiation^20,49,59^. BMP-associated *Smad1/5* are enriched in the AEP TRN while Tgfβ-associated *Smad2/3* are enriched in the AT1 TRN, further supporting the observation that the AT1tr state is a differentiation intermediate between AEP and AT1. The top predicted inhibitory factor in the AT1tr TRN is *Bach1*, a known modifier of cellular metabolic state^65,66^. Bach1 represses expression of many genes predicted to be activated by *Mef2* factors, which function as regulators of cellular fate and cytoskeletal organization in diverse cell types^67^. Mef2c was recently identified as a key potential regulator of AT2 to AT1 transitions in two-dimensional culture of human lung epithelia^68^, suggesting balance of Mef2 activation and Bach1 repression in AT1 differentiation.

### Loss of Nkx2-1 activity defines the Krt8^+^ transitional cell TRN

Comparing the AT1tr and Krt8^+^ cell states highlighted a striking absence of predicted Nkx2-1 activity in Krt8^+^ cells. TRNs from all other epithelial cell states in AEP-O included Nkx2-1. Recent epigenomic profiling of the activity of Nkx2-1 during AT2 to AT1 transitions demonstrated Nkx2-1 occupancy at different genomic regions in each AT2 vs AT1 cells^25^, suggesting a role for Nkx2-1 dis-engagement and re-engagement in the genome during differentiation. Nkx2-1 expression is lowest in stressed transitional epithelial cells at the time when Krt8 expression is highest during *in vivo* lung regeneration^22^. Together, these observations supported the hypothesis that reduction in Nkx2-1 activity could directly promote transition to the stressed Krt8^+^ transitional state.

To test this concept, we developed an approach to genetically manipulate AEPs during formation of AEP-O (Figure 5). Using an AAV6.2FF-Cre^69^, which has recently been described as a high-fidelity reagent for genetic manipulation of AT2 cells *in vivo*^*70*^, we infected AEPs harboring a R26R-lox-stop-lox-EYFP allele^71^ (from R26R^EYFP^ mice) immediately after FACS sorting (Figure 5A). AAV6.2FF-Cre efficiently targeted AEPs which produced morphologically complex organoids (Figure 5B) expressing the EYFP lineage label (Figure 5C). Titration experiments indicated that infection at a multiplicity of infection (MOI) of 1000 was sufficient to induce significant recombination and label the majority of AEP-O (Figure 5D); higher MOI mildly increased targeting, but at the expense of reduction in colony formation. Multiple biological replicates confirmed that MOI of 1000 led to targeting of approximately 60% of organoids with no change in colony formation efficiency or size of organoids (Figure 5E); whole mount IHC confirmed EYFP expression with no reduction in internal complexity of AEP-O (Figure 5F). These results indicated that AAV6.2FF-Cre was capable of efficiently targeting AEPs *in vitro* for genetic manipulation without perturbing the AEP-O system and confirmed the clonal nature of AEP-derived organoids.

**Figure 5.**
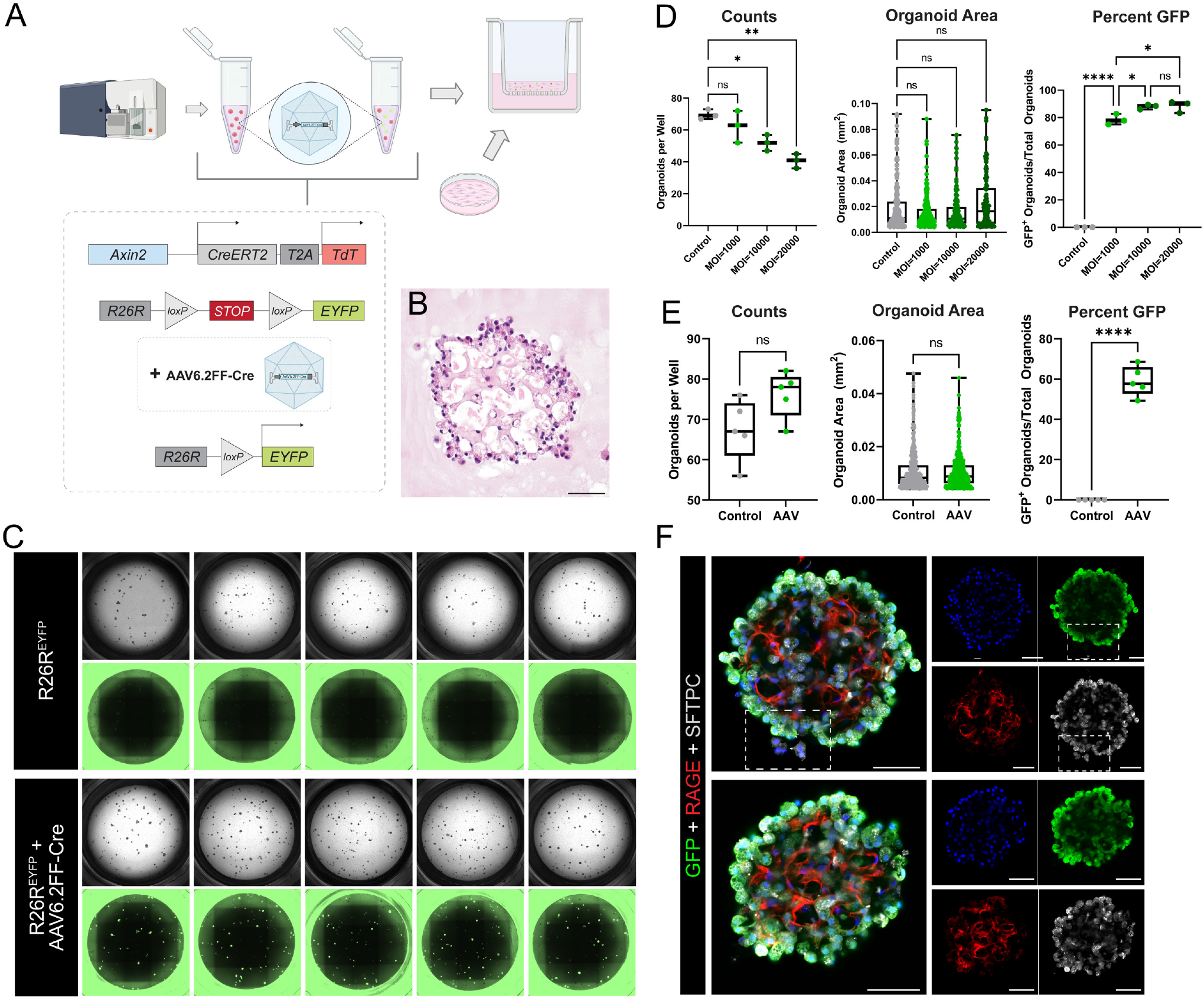
*In vitro* gene editing of AEP-derived alveolar organoids via AAV6.2FF-Cre. AAV6.2FF-Cre experimental set-up. Live/CD31^-^/CD45^-^/CD326^+^(EpCAM^+^)/TdTomato^+^(Axin2^+^) cells (AEPs) sorted from mice with the R26R^EYFP^ allele (Axin2^creERT2-tDT^; R26R^EYFP^) were treated with AAV6.2FF-Cre and plated with wild-type fibroblasts. (B) H&E of 5 μm sections of FFPE day 29 AAV6.2FF-Cre-treated organoids, exhibiting morphology and structural complexity similar to untreated/control organoids (Figure 1B). (C) Whole-well brightfield and GFP images of day 29 organoids (untreated vs. AAV6.2FF-Cre-treated) at MOI=1000. (D) Comparison of cells treated with an MOI of 1000, 10000, 20000. Quantification of day 32 organoids (n=3 wells per condition) showing that an MOI of 1000 causes recombination in organoids without significant effects on colony forming efficiency (CFE). (E) Quantification (n=5 wells per condition) showing that an MOI=1000 induces significant levels of recombination without significant effects on organoid number or size. (F) Whole-mount immunofluorescence of day 32 AAV6.2FF-Cre-treated AEP-derived organoids (same experimental set-up as Figure 2). White box highlighting untargeted epithelial cells (YFP^-^/GFP^-^) next to a targeted (YFP^+^/GFP^+^) organoid in the same well, supporting clonal expansion of AAV6.2FF-Cre-treated cells. (*ns = p > 0*.*05; *P =0*.*05, **P =0*.*01, ***P =0*.*001, and ****P =0*.*0001*). Note: EYFP was stained for using anti-GFP antibodies and imaged (whole well images) using GFP filter cubes. [*Scale bars = 50 μm*]. *AAV = Adeno-Associated Virus; MOI = Multiplicity of Infection*

### Nkx2-1 deficient AEPs transition to the Krt8^+^ stressed transitional cell state

To test the hypothesis that Nkx2-1 deficiency contributed to the Krt8+ state transition, we applied AAV6.2FF-Cre to AEPs from Axin2^CreERT2-tdT^ x R26R^EYFP^ x Nkx2-1^flox/flox^ animals immediately after sorting. This generated Nkx2-1 knockout AEPs which were used to initiate organoid formation (Figure 6A). Morphology in EFYP^+^ (Nkx2-1^KO^) organoids was noticeably different from EYFP^-^ organoids in the same well, and EFYP^+^ organoids were substantially larger by day 28 of culture (Figure 6B). Because AAV6.2FF-Cre only targeted ∼60% of organoids per well, we were able to directly compare EFYP^+^ and EFYP^-^ organoids grown in the same well to assess the impact of Nkx2-1 knockout in AEP-O. As expected, we noted robust Nkx2-1 protein expression in EFYP^-^ organoids (Figure 6C-E), and complete loss of Nkx2-1 protein in EFYP^+^ organoids. Nkx2-1^KO^ AEPs lost expression of AT2 markers, including Sftpc, with associated increased expression of E-cadherin (Cdh1) and change in cell shape and organoid morphology (Figure 6F-H). Diverse morphological types were visible in Nkx2-1^KO^ AEP-O, with loss of alveolar-like cavities, prominence of one or a small number of large cavities full of debris, and pseudostratified epithelial lining with some organoids exhibiting a glandular appearance (Figure 6F,I). These structures were reminiscent of other endoderm-derived organs, including the esophagus, stomach, and intestine. Nkx2-1 knockout in differentiated distal lung lineages has been associated with expression of foregut endoderm genes^25,72-74^, so we examined protein expression of a large panel of proximal lung and foregut endoderm markers including Sox2, Sox9, Cdx2, Gata4, and Pdx1. No substantial expression of any of these markers was detectable, suggesting Nkx2-1 KO AEP-O epithelium did not adopt foregut endodermal fate from proximal lung or GI organs (Figure S6 A-Q).

**Figure 6.**
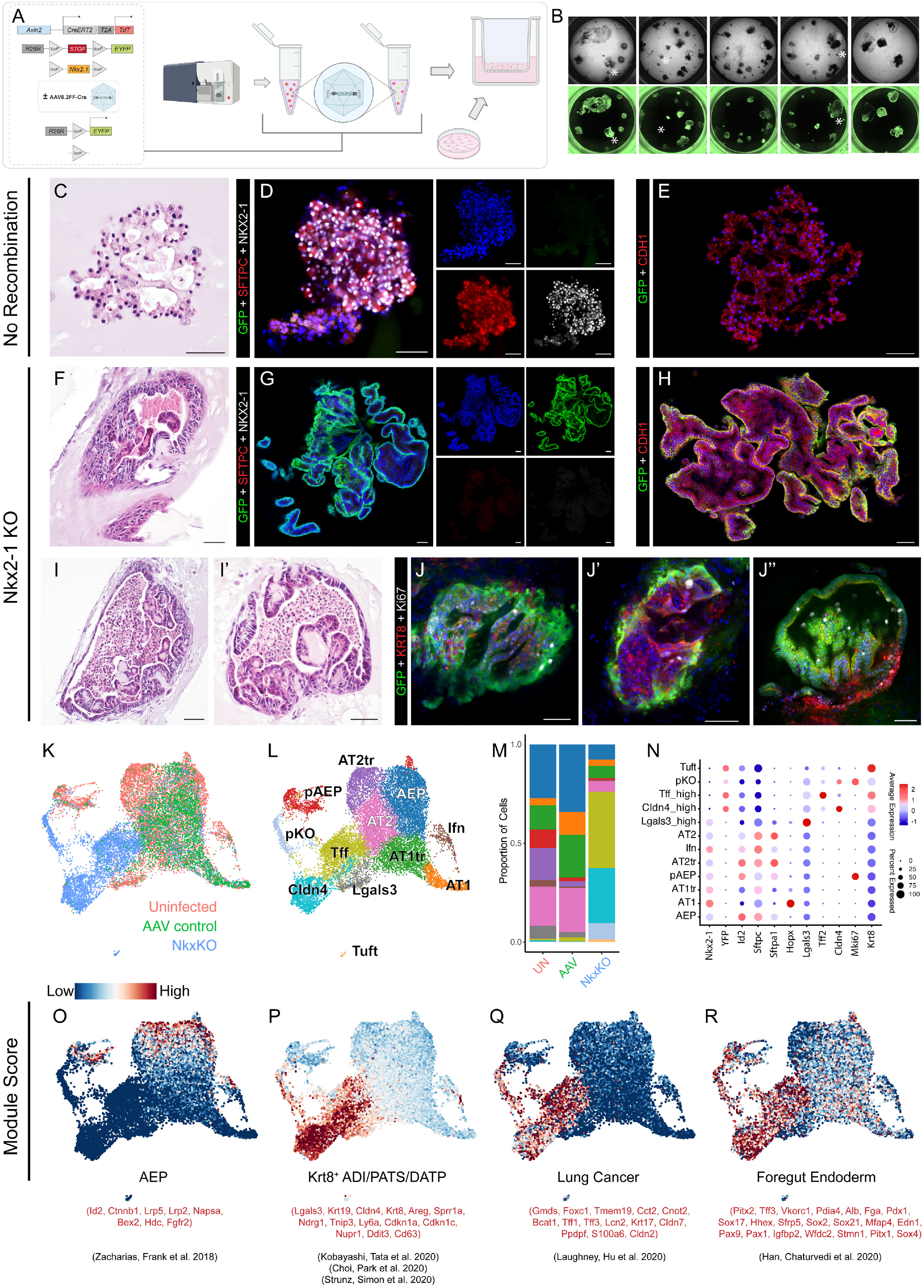
*In vitro* Nkx2-1 KO of AEP-derived alveolar organoids drives irreversible transition to a Krt8^+^ stressed transitional state. (A) AAV6.2FF-Cre experimental set-up. Live/CD31^-^/CD45^-^/CD326^+^(EpCAM^+^)/TdTomato^+^(Axin2^+^) cells (AEPs) sorted from Axin2^creERT2-tDT^; R26R^EYFP^ mice and Axin2^creERT2-tDT^; R26R^EYFP^ ; Nkx2-1^fl/fl^ mice were treated with AAV6.2FF-Cre and plated with wild-type fibroblasts. (B) Comparison of brightfield and GFP whole-well images of organoids grown from control (AAV6.2FF-Cre-treated sorted R26R^EYFP^ AEPs) and Nkx2-1 KO AEPs (AAV6.2FF-Cre-treated sorted R26R^EYFP^; Nkx2-1^fl/fl^ AEPs) at day 28 of culture. Control (non-GFP) organoids with normal morphology are marked with white asterisk. (C-J) H&E and immunofluorescence images of R26R^EYFP^; Nkx2-1^fl/fl^ AEP-derived organoids that did (F-J) or did not (C-E) undergo recombination via AAV6.2FF-Cre. (C-E) Non-recombined organoids (D) express SPC (red) and Nkx2-1 (white), but do not express the YFP lineage label (green), whereas (G) recombined organoids do not express SPC or Nkx2-1 but do express the YFP lineage label. Non-recombined (E) and recombined (H) organoids maintain epithelial identify expressing CDH1. Nkx2-1^KO^ organoids express KRT8 and many proliferate and express Ki67 at late as day 40 of culture (J-J’’). (K-R) Integrated scRNAseq comparing epithelial cells from day 28 control organoids (Uninfected), AAV6.2FF-Cre-treated control organoids (AAV control), and AAV6.2FF-Cre-treated Nkx2-1 KO organoids (Nkx2-1^KO^). Nkx2-1^KO^ cells cluster separately from Uninfected and AAV control cells near Krt8^+^ cells (K-L), which make up a majority of cells in the Nkx2-1^KO^ condition (M). Marker genes for normal alveolar epithelium are lost and novel markers gained (N) in Nkx2-1^KO^. (O-R) Module scoring using published gene sets for AEPs (O), Krt8/PATS/DATP/ADI cells (P), lung cancer cells (Q), and foregut endoderm (R). Compare to Figure S7 for detailed marker gene analysis.

To clarify Nkx2-1^KO^ AEP-O composition, we turned to unbiased profiling. We performed scRNAseq in Nkx2-1^KO^ AEP-O at 28d of culture and compared the expression profile of Nkx2-1^KO^ cells with Nkx2-1 expressing cells in our WT organoid scRNAseq time series. We added a control condition of AAV6.2FF-Cre treatment in AEP-O from Axin2^CreERT2-tdT^ x R26R^EYFP^ (as in Figure 5), to rule out any AAV6.2FF-Cre-specific effects. We then integrated scRNAseq expression data from Nkx2-1^KO^ and Nkx2-1^+/+^ AEP-O and compared gene expression profiles (Figure 6K-R, S7). Consistent with IHC, Nkx2-1 and EYFP RNA expression were mutually exclusive. Nkx2-1 was undetectable in EYFP^+^ epithelial cells at the RNA level (Figure 6N). EYFP^+^ Nkx2-1^KO^ cells form multiple distinct clusters separated from control cell types (Figure 6K). Nkx2-1^KO^ cells clustered near WT Krt8^+^ transitional epithelial cells (Figure 6L) and expressed high levels of transcripts enriched in the of Krt8^+^/PATS/DATP/ADI state including *Cldn4, Tff2*, and intermediate levels of *Lgals3* (Figure 6N, S7Q-T). A distinct proliferative cluster was present by scRNAseq among *Cldn4*-high Nkx2-1^KO^ cells (Figure 6K). Ki67 expression in Krt8^+^ Nkx2-1^KO^ AEP-O demonstrated ongoing proliferation at 40d of culture despite large organoid size (Figure 6J).

Given the close association of Nkx2-1^KO^ epithelial cells to the Krt8^+^ state in Nkx2-1 WT AEP-O, we used Seurat module scoring to compare Nkx2-1^KO^ and WT cells. Cells in Nkx2-1^KO^ organoids lost AEP-associated gene expression (Figure 6O) while activating cell stress markers associated with Krt8^+^ cells in WT AEP-O. Cells from Nkx2-1^KO^ AEP-O were highly enriched for gene sets associated with human lung adenocarcinoma^75^, concordant with previous findings implicating Nkx2-1 loss in the pathogenesis of lung cancer^72,74^. While we did not detect protein expression of non-lung foregut endoderm markers by IHC, Nkx2-1 KO epithelial cells did express low levels of non-lung endodermal genes at the RNA level^76^, consistent with loss of the instructive activity of Nkx2-1 in constraining lung fate. Taken together, these data support the concept that Nkx2-1 loss in AEPs led to acquisition of the Krt8^+^/PATS/DATP/ADI cell state, validating the prediction of the Krt8^+^ cell state TRN analysis.

### Deletion of Nkx2-1 in AEPs in vivo caused rapid, spontaneous conversion to a proliferative Krt8^+^ transitional state

To further evaluate the hypothesis that Nkx2-1 loss is sufficient to induce the Krt8^+^ cell state in AEPs, we performed *in vivo* lineage tracing of Nkx2-1^KO^ AEPs in adult mice at homeostasis. While Axin2^CreERT2^ functions as an effective lineage tracing reagent for lung epithelial cells after high dose tamoxifen treatment^3,18^, the relative CreERT2 recombination inefficiency with this line prevented full deletion of floxed alleles in lineage labeled cells in adult lung epithelium at high doses of tamoxifen (data not shown). We therefore used the recently reported epithelial-specific, AEP-enriched^48^ Tfcp2l1^CreERT2^ mouse line^77^ to generate Tfcp2l1^CreERT2^ x R26R^EYFP^ x Nkx2-1^flox/flox^ animals, enabling Nkx2-1 knockout in AEPs during adult alveolar homeostasis (Figure 7A). In Nkx2-1^wt/wt^ animals, we detected Tfcp2l1^CreERT2^ lineage-label in solitary AT2 cells (SFTPC^+^/NKX2-1^+^) scattered throughout individual alveoli after administration of tamoxifen as expected for an AEP-enriched Cre line (Figure 7B-E).

**Figure 7.**
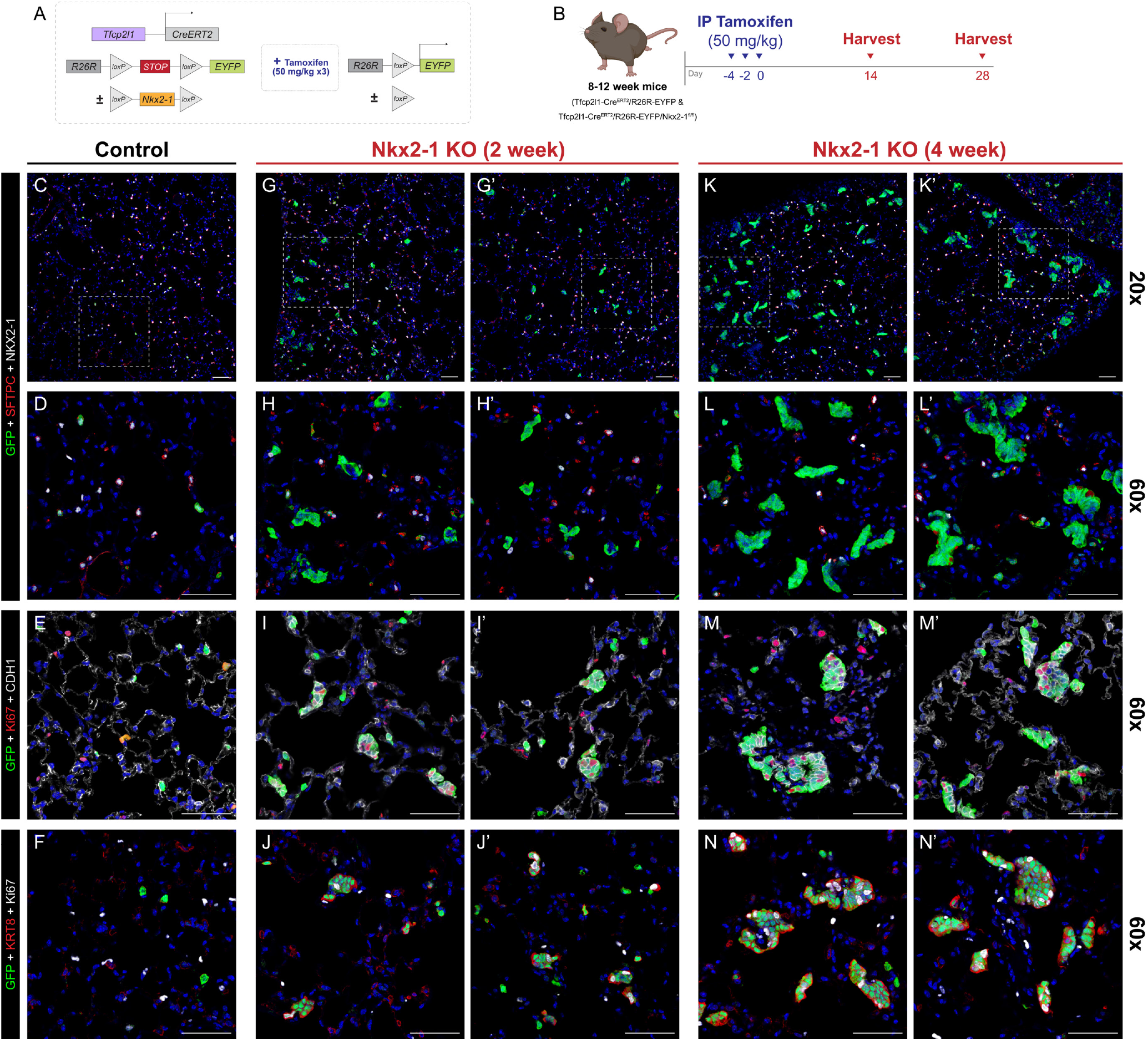
Genetic deletion of Nkx2-1 in vivo leads to loss of distal lung fate and acquisition of PATS/Krt8^+^ state. (A-B) Experimental design of *in vivo* genetic ablation of Nkx2-1 in AEPs. Mouse genetic construct (A) and experimental treatment plan and schematic (B). 8-12-week Tfcp2l1-CreERT2; R26R^EYFP^ (C-F) and Tfcp2l1-CreERT2; R26R^EYFP^; Nkx2-1^fl/fl^ (G-N) were treated with three doses of IP tamoxifen (50 mg/kg) and harvested at 2 to 4 weeks post-treatment; control is from 2-week timepoint. (C) Control (Tfcp2l1-CreERT2; R26R^EYFP^) mice exhibited YFP induction in a subset of AT2 cells (SPC^+^ [red]/Nkx2-1^+^ [white]) with normal histological characteristics. (G-H, K-L) Nkx2-1 KO (Tfcp2l1-CreERT2; R26R^EYFP^; Nkx2-1^fl/fl^) mice exhibited clustered YFP^+^ proliferative clones negative for AT2 cell markers (SPC^-^ [red]/Nkx2-1^-^ [white], H, L), with acquisition of Ki67 and Krt8 expression (I-J, M-N). Progressive clonal enlargement by 4 weeks post treatment (K) disrupts normal lung morphology, with continued growth and proliferation (all scale bars = 50 μm).

At 2 weeks following Nkx2-1 deletion in AEPs, we detected multi-cellular clones of lineage labeled EYFP^+^, Nkx2-1^KO^ epithelial cells throughout the lung (Figure 7F-I). Nkx2-1^KO^ Tfcp2l1-lineage cells lost expression of AT2 markers including Sftpc, changed shape, and acquired an E-cadherin high, Krt8-expressing, proliferative state (Figure 7J-M), consistent with changes seen in Nkx2-1^KO^ organoids. At 4 weeks of age, these Krt8^+^ clones had grown substantially, with persistent shape change and high-level proliferation (Figure 7N-U). Disruption of alveolar morphology was present in some areas near larger clones (Figure 7M). Taken together, these findings confirm that, as in AEP-O, Nkx2-1 loss in AEPs *in vivo* drives acquisition of the Krt8^+^/PATS/DATP/ADI molecular state, with spontaneous proliferative growth and disruption of alveolar architecture (Figure 8).

**Figure 8.**
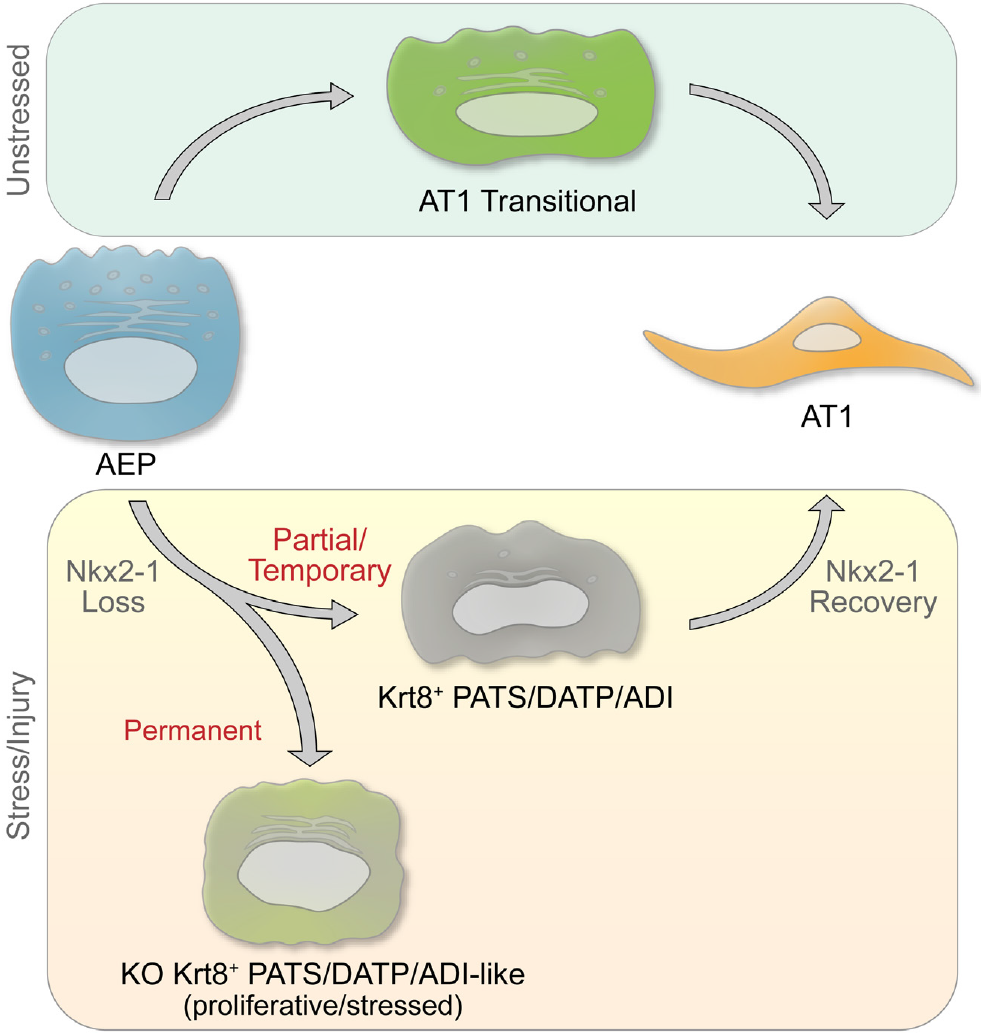
Model of Nkx2-1 activity in controlling progenitor and transitional cell state. AEP can differentiate to AT1 cells via either the AT1tr or Krt8^+^ states during homeostasis, with Nkx2-1 release from AT2 genes during transition through a Krt8^+^ state^25^. Nkx2-1 activity and expression are lowest in Krt8^+^ cells^22^, and Nkx2-1 must re-engage chromatin to complete AT1 transition from the Krt8^+^ state^25^. Permanent Nkx2-1 loss in AEPs causes transition to proliferative, stressed, Krt8^+^-like state characterized by unconstrained growth *in vitro* and *in vivo*. Nkx2-1 is therefore required for the progenitor activity of AEPs and supports the transition to AT1 cells by Krt8^+^ cells.

## Discussion

In summary, combined scRNAseq and scATACseq of AEP-derived alveolar organoids allowed definition of regulatory networks along multiple differentiation trajectories of lung epithelial progenitor cells toward differentiated alveolar epithelium. Transcriptomic and epigenomic data demonstrated two distinct AEP to AT1 differentiation trajectories, one through a known stressed transitional cell state and another through a state defined by signaling integration and transcriptional regulators of AT1 cell gene programs. A major difference in the regulation of these two cell states was activity of Nkx2-1. Nkx2-1 expression in AEPs is required to maintain the AEP state, and loss of Nkx2-1 activity is sufficient for AEPs to enter the Krt8^+^ stressed transitional state. Nxk2-1 therefore plays a crucial, previously unrecognized role in the maintenance of progenitor function in the adult lung. These findings were enabled by the characteristics of the AEP-O system, allowing close interrogation of progenitor dynamics in a model with similar cellular and morphological complexity to alveolar regeneration *in vivo*.

### Under-recognized key roles of developmental factors in diverse homeostatic and regenerative biology

Nkx2-1 loss is sufficient to cause transition of AEPs to a proliferative stressed transitional state, a finding which emphasizes the need for active maintenance of adult AT2 alveolar progenitor capacity. At a basic level, the centrality of Nkx2-1 in lung progenitors is not surprising, as prior reports have demonstrated a requirement for Nkx2-1 expression in maintenance of lung epithelial fate in adult differentiated cells^25,54,72-74^. Loss of one or more alleles of Nkx2-1 is a common mutation found in lung adenocarcinoma^78^. At a deeper level, however, it is quite provocative that the loss of a single transcription factor in a facultative progenitor lineage is sufficient to drive not only a fate transition but the conversion of a poised quiescent lineage to unconstrained proliferative growth. It is increasingly understood that transcription factors exert influence on gene expression through changes in chromatin state at regulatory elements, especially so called “pioneer factors". Nkx family factors are known to function as pioneer transcription factors in diverse contexts^79^, and pioneer factors catalyze changes in chromatin structure that maintain epigenetic stability^80^. There is a clear need to define with precision the mechanisms underlying the dynamic Nkx2-1 activity in chromatin reorganization^25^. Important first steps include addressing the interactions of common chromatin regulatory complexes with known lung lineage factors including Nkx2-1, Foxa, Gata, and Hopx, and to carefully evaluate changes in and maintenance of the epigenomic state and chromatin topology across time during the lifespan of the lung.

### Evaluation of epigenomic state resolves complexity underlying divergent cell states

The identification of Nkx2-1 as a regulator of AEP progenitor state relied on the ability to distinguish similar cell states in a dynamic system. To that end, we utilized a clonal system with known input/initial cell state obtained by sorting enriched AEPs, providing a clear ability to identify the starting point of differentiation. Identification of initial root cell in trajectory, pseudotime, and RNA velocity analyses can be challenging and the shortfalls can be mitigated by controlling input^44,81^. Knowing the input cell state of AEP-O added confidence when the unbiased methods correctly identified AEPs or proliferating AEPs as the initial state. Together, these factors enabled downstream identification of AT1 differentiation trajectories through both the AT1tr and Krt8^+^ states. Addition of chromatin state provided a more refined signature for AEPs, allowing better identification of AEPs within the AT2 population, which has been challenging based on scRNA transcriptome alone^17,37^. Together, scRNA and scATACseq provided the resolution necessary to derive TRN inference, leading to Nkx2-1. Beyond the biological impact of Nkx2-1 deletion in AEPs, the marked phenotype provides a clear proof of principle that addition of chromatin topology and regulatory network inference can identify unexpected regulators of lung biology. There are several other testable predictions of the AEP-O TRN, which will need to be validated individually. The reproducibility and flexibility of an organoid system will enable these studies, as reagents such as AAV6.2FF provide a platform for rapid screening of novel regulators of alveolar regeneration.

### Transitional cells at the interface of cell stress, regeneration, and disease

The marked differences between AT1tr and Krt8^+^ transitional cells further the complexity of choosing cellular targets for lung regeneration. Krt8^+^ stressed transitional cells have been described in diverse models, are found in mouse and human, are increased in several disease states, and are readily identifiable based on high level expression of enriched markers^82^. However, our data suggest that permanent acquisition of a stressed transitional state, such as seen in Nkx2-1 KO organoids and Nkx2-1 KO AEPs *in vivo*, may drive aberrant proliferation, expression of lung cancer programs, and loss of lung identity. Progressive acquisition of this cell state may therefore be deleterious; even if most cells pass through or ‘recover’ from this stressed transitional state, accumulation of ‘stuck’ transitional cells may represent a risk factor for development of lung disease. Our data suggest two potential avenues to move forward in identifying actionable therapeutic strategies. First, to identify factors, such as Nkx2-1, required for progression of stressed transitional cells toward AT1 fate to promote progression or “rescue” stuck cells. Second, to determine mechanisms to promote progenitor cells to pass through the less stressed AT1tr transitional state as they differentiate. Future studies designed to address the determinants of these transition states during lung injury and modifiers of progression during regeneration and disease are therefore of high priority. Given the accumulation of stressed transitional cells in fibrosis^19^ and relative transience in acute regeneration^22^, both strategies are likely to be useful in different clinical contexts.

### AEP-Os are a high-fidelity ex vivo model of alveolar epithelial regeneration

Restoration of a gas exchange surface through repair or replacement of injured alveoli is the central process needed to promote therapeutic regeneration. Here, we show that AEP-derived lung organoids recapitulate the major stages of the epithelial portion of the alveolar regenerative process, modeling progenitor cell expansion, alveolar epithelial differentiation, and formation of alveolar-like cavities with properly polarized and organized epithelium. Nkx2-1 deletion caused concordant changes both in organoids and *in vivo*, providing proof of principle that AEP-O model key aspects of adult alveolar biology and alveolar regeneration. While we characterized AEP-O only from mice in this study, prior reports have demonstrated capacity for human AT2 progenitor culture^3,7,11,83^, and advancing human primary alveolar progenitor culture is a high priority for therapeutic development in tandem with continued progress in stem cell-based organoid technology.

Controversy exists regarding the similarity of alveolar regeneration and alveolar development^84^; Some have argued that regeneration is fundamentally different due to the altered milieu of the injured alveolus, or even suggested that pathological remodeling rather than functional regeneration is the end state of a significant portion of lung injury^85^. Our findings argue that extensive regenerative potential is encoded in the AEP epigenomic state, driving a distinct process to rebuild alveoli. AEPs contain the required information to undergo progenitor self-renewal, multilineage differentiation, and complex morphogenesis in the presence of a minimal signaling niche. This challenge bears striking resemblance to the injured lung, where epithelial, mesenchymal, endothelial, immune lineages and the underlying matrix environment are all altered by pathogens^86^. AEP-driven cavity formation occurs in the absence of mechanical contribution from myofibroblasts, quite different from during alveologenesis when the mechanical activity of myofibroblasts is required for formation of alveoli^87,88^. Our data therefore emphasize the difference between alveolar development and alveolar regeneration and suggest that AEP-O provide unique benefits to study regenerative biology specifically. Continued advancement of primary lung organoids with diverse cell compositions will provide models which balance between fidelity and reproducibility, enabling and accelerating progress in defining molecular mechanisms of lung regeneration.

## Acknowledgements

The authors would like to thank Cheng-Lun Na (CCHMC) for his expertise and assistance with electron microscopy, and Tara Rindler for preparation of the AAV6.2FF-Cre. The authors would also like to thank the CCHMC Confocal Imaging Core (CIC) (especially Matt Kofron and Sarah McLeod), the CCHMC Single Cell Genomics Core (SCGC) (especially Kelly Rangel and Shawn Smith), the CCHMC Division of Bioinformatics (for providing/maintaining the high-performance computational cluster), the CCHMC DNA Sequencing Core, and the CCHMC Research Flow Cytometry Core (RFCC) in the Division of Rheumatology. Select figure schematics were created using BioRender.com.

## Author Contributions

Experimental Design – AT, PK, WJZ

Data Acquisition – AT, PK

Data Analysis – AT, JS, MK, JAW, ERM, DS, WJZ

Provision of Critical Reagents and Methods Expertise – JPB, MK

Writing of first draft of paper – AT, WJZ

Editing and critical revision – AT, PK, JS, MK, JAW, JPB, ERM, DS, WJZ

Acquisition of funding support – WJZ, ERM

## Funding

AT was supported by HL007752 (NIH), ERM by AI150748 (NIH), and WJZ by HL140178 (NIH), AI150748 (NIH), and the CCHMC Procter Scholar Award.

## Competing interests

The authors declare no competing interests for the presented work (including patents, financial holdings, advisory positions, or other interests).

## Data/Code/Protocol/Material Availability

All raw data has been uploaded to GEO database and are available directly by request to the contact PI. All data is also undergoing upload to the LungMAP web portal (https://lungmap.net/explore-data/) for full visualization and individual exploration.

## Materials and Methods

To optimize important aspects of the AEP-O culture, we undertook extensive reagent testing and standardization. The primary components of epithelial organoid co-culture assays are 1) epithelial cells; 2) supportive cells, if any; 3) matrix for three-dimensional suspension and growth; 4) media and media additives; and 5) growth surface (i.e., Transwell filter). We tested each of these components iteratively. As previously reported, AEPs form more and larger organoids than unselected AT2 cells^3^, so we focused on using FACS-sorted AEPs derived from Axin2^CreERT2-tdT^ mice as the epithelial starting fraction. To control for observed variability in organoid morphology and composition from different types of supportive mesenchymal cells, we used large, consistent preparations of primary lung fibroblasts obtained by selective adhesion from P28 wild type C57BL/6 mice at passage 3-4. We then lot tested 4 available commercial matrices, all with composition similar to Matrigel (Corning), and found that the best lots of Matrigel support organoid growth with 2-3-fold increased CFE compared to other Matrigel lots and competing products; we therefore confined our studies to a single lot of Matrigel, and all data herein uses this standardized reagent. We cultured 5,000 FACS-purified AEPs and 50,000 lung fibroblasts in each well on a Transwell filter in a 1:1 ratio of Matrigel and media (SAGM with 5% FBS and limited additives); we took this approach minimize exogenous signaling modulators in the media and allow evaluation of the supportive capacity of the mesenchymal fraction. To evaluate the growth of these organoids more fully, we adapted methods from iPSC culture for whole mount immunohistochemistry of organoids^28^.

### Ethical Compliance and Animals

All animal studies were conducted under the guidance and supervision of the Cincinnati Children’s Hospital Medical Center (CCHMC) Institutional Animal Care and Use Committee (IACUC) in accordance with CCHMC regulatory and biosafety protocols. Mouse lines used included: C57BL/6J mice (Jackson Strain #000664), PDGFRα^EGFP^ (B6.129S4-PDGFRa^tm11(EGFP)Sor^/J; Jackson Strain #007669) Axin2^creERT2-TdT^ (a gift from Edward Morrisey), Tfcp2l1^CreERT2^ (B6;129S-*Tfcp2l1*^*tm1*.*1(cre/ERT2)Ovi*^*/J; Jackson Strain* #028732), Nkx2-1^fl/fl^ (a gift from Shioko Kimura), and R26R^EYFP^ (B6.129X1-Gt(ROSA)26Sor^tm1(EYFP)Cos^/J; Jackson Strain #006148). All experiments for both organoids and *in vivo* lineage tracing included both male and female mice. For Cre recombinase induction in mouse models, 8–12-week-old mice were treated intraperitoneally (IP) with Tamoxifen (Sigma, T5648; dissolved in ethanol and resuspended in corn oil) at a dose of 50 mg/kg, one or three times (every other day), at the experimental timepoints indicated previously.

### Mouse Lung Harvest

Mice were anesthetized via IP injection of a lethal dose of ketamine and xylazine followed by confirmation of euthanasia via cervical dislocation and thoracotomy. The chest cavity was opened to expose the heart and lungs. The right ventricle was perfused with 5-10 mL of cold PBS (Gibco, 10010-023) to clear blood from the lungs. For tissue dissociation for organoids, lungs were removed and placed in cold PBS on ice. For tissue fixation for histology and immunofluorescence, the trachea was cannulated and lungs were inflated to a pressure of 30 cm H2O using 4% paraformaldehyde (PFA). Inflated lungs were immersed in a conical of 4% PFA, then left on a rocker at 4°C overnight.

#### Processing Fixed Lung Tissue for Histology & Immunofluorescence

The day following inflation, fixed lung tissue was trimmed and placed in cassettes. The cassettes were washed (15 minutes each) 3x in DEPC-treated PBS, 1x in DEPC-treated 30% ethanol, 1x in DEPC-treated 50% ethanol, and 3x in DEPC-treated 70% ethanol. Following a standardized overnight automated processing protocol (Thermo Scientific, Excelsior ES), the samples were embedded in paraffin. Samples were sectioned at a thickness of 5 μm. Paraffin sections were incubated at 65°C for two hours, deparaffinized in xylene (3x for 10 minutes), rehydrated through an ethanol gradient, and standard H&E staining was performed. Slides were mounted with Permount Mounting Medium (Electron Microscopy Sciences, 17986-05) and cover slipped with #1.5 Gold Seal 3419 Cover Glass (Electron Microscopy Sciences, 63790-01). Immunofluorescence was on paraffin sections was performed as previously described^3^. Briefly, following deparaffinization, rehydration, and sodium citate antigen retrieval (10 mM, pH 6.0), and blocking, immunofluorescence was performed on paraffin sections using antibodies in Supplemental Table 1b and the following reagents: ImmPRESS(r) HRP Horse Anti-Rabbit IgG Polymer Detection Kit (Vector Labs, MP-7401-50), ImmPRESS(r) HRP Horse Anti-Goat IgG Polymer Detection Kit (Vector Labs, MP-7405-50), and ImmPRESS(r) HRP Goat Anti-Rat IgG, Mouse adsorbed Polymer Detection Kit (Vector Labs, MP-7444-15). Following application of TSA fluorophores (listed in Supplemental Table 1b; 1:100), sections were stained with DAPI (Invitrogen, D1306; 1:1000) and mounted using Prolong Gold antifade mounting medium (Invitrogen, P36930).

### Mouse Lung Digestion and Single Cell Suspension

Clonal mouse alveolar epithelial progenitor (AEP)-based alveolar organoids were generated as previously described^3^ with minor modifications. Briefly, following harvest, lungs were removed from ice cold PBS and non-pulmonary tissue and gross airways were removed via manual dissection, and lung tissue was finely chopped and transferred to a GentleMACS C tube (Miltenyi Biotec, 130-093-237) (tissue from one mouse per C tube) containing 5 mL of digestion buffer [composed of 9 mL of phosphate-buffered saline (PBS; Gibco, 10010-023) combined with 1 mL of Dispase (stock: 50 U/mL; final concentration: 5 U/mL, Corning, 354235), 50 μL of DNase (stock: 5 mg/mL; final concentration: 0.025 mg/mL or 50 U/ml, GoldBio, D-301), and 100 μL of Collagenase Type I (stock: 48,000 U/mL; final concentration of 480 U/mL, Gibco, 17100-017)]. C tubes were placed on a gentleMACS Octo Dissociator with Heaters (Miltenyi Biotec, 130-096-427) and the following protocols were run: “*m_lung_01_02*” (36 seconds) twice, “*37C_m_LIDK_1*” (36 minutes 12 seconds) once, and “*m_lung_01_02*” (36 seconds) once. Samples were passed through a 70 μm filter (Greiner Bio-One, 542070) and centrifuged at 500g for 5 minutes at 4°C. Following removal of the supernatant, 5 mL of RBC Lysis Buffer (Invitrogen, 00-4333-57) was added and incubated for 5 minutes. All centrifugation steps with this single cell suspension were performed at 500g for 5 minutes at 4°C for the following procedures.

### Fibroblast Stock Preparation (with PDGFRα^EGFP^ fibroblasts)/Media

For generation of fibroblast stocks, 4-week C57BL/6J mice and 4-week PDGFRα^EGFP^ mouse lungs were harvested, digested, and processed as described above. Following centrifugation, cells were washed 3x with MACS Buffer (autoMACS Rinsing Solution [Miltenyi Biotec, 130-091-222] with MACS BSA Stock Solution [Miltenyi Biotec, 130-091-376]). After removing supernatant from final wash, the cell pellet was resuspended in 10 mL fibroblast medium (DMEM/F-12 [Gibco, 11320-033], Antibiotic-Antimycotic [Gibco, 15240-062, final concentration 1x], and Heat Inactivated Fetal Bovine Serum [Corning, 35-011-CV, final concentration 10%]) and plated on a 10 cm tissue culture plate (approximately 1 mouse per plate). Non-adherent cells were removed via media change 2-12 hours post-plating.

Cells were passaged at 80% confluency to P3. For passaging, media was removed from each plate and cells were washed with 5 mL of DPBS (Gibco, 14190-094). 3 mL of 0.25% Trypsin-EDTA (Gibco, 25200-056) was added and plates were incubated at 37°C for 7 minutes. 5 mL of fibroblast medium was added to each plate, pipetted to dissociate cells, and transferred a 15 mL conical tube. Cells were centrifuged at 500g for 5 minutes at 4°C, supernatant was removed, and cell pellet was resuspended in 2 mL fibroblast medium/per plate (split 1:2 or 1:3) and transferred to plates containing 6 mL fibroblast medium.

Once confluent at P3, cells were washed, trypsinized, centrifuged as above, and resuspended in 1 mL of freezing medium (90% FBS, 10% DMSO) (one plate per cryovial) and transferred to Mr. Frosty Cryogenic Freezing Container (Nalgene, 5100-0001) filled with isopropyl alcohol, which were placed in a -80°C freezer overnight, before samples were moved to long-term liquid nitrogen storage.

For use of frozen fibroblast stocks in organoids, 48 hours prior to use in organoids, cells were rapidly thawed at 37°C and resuspended in 10 mL fibroblast medium in a 15 mL conical. Cells were centrifuged at 500g for 5 minutes at 4°C, supernatant was removed, and cell pellet was resuspended in 2 mL fibroblast medium and transferred to a 10 cm tissue culture plate containing 6 mL of fibroblast medium. Fibroblasts used for organoids were washed, trypsinized, and resuspended before counting.

### Processing for Organoids/FACS/Cell Sorting

Single cell suspensions were obtained as above, and cells were resuspended in 5 mL MACS Buffer (autoMACS Rinsing Solution [Miltenyi Biotec, 130-091-222] with MACS BSA Stock Solution [Miltenyi Biotec, 130-091-376]) and passed through a 40 μm filter (Greiner Bio-One, 542040). Cells were centrifuged, the supernatant was removed, and the cell pellet was resuspended in Fc Receptor Binding Inhibitor Polyclonal Antibody (Invitrogen, 14-9161-73) diluted 1:100 in MACS buffer and incubated for 10 minutes at room temperature. Following centrifugation, cells were resuspended in a mixture of the following antibodies diluted 1:100 in MACS buffer and incubated for 10 minutes protected from light: CD31 (PECAM-1; Monoclonal Antibody [390], eFluor 450) (Invitrogen, 48-0311-82), CD45 (Monoclonal Antibody [30-F11], eFluor 450) (Invitrogen, 48-0451-82), CD326 (EpCAM; Monoclonal Antibody [G8.8], APC) (Invitrogen, 17-5791-82). Cells were washed 1x with 1-5 mL of MACS buffer, and resuspended in Fixable Viability Dye eFluor 780 (Invitrogen, 65-0865-14) diluted 1:1000 in MACS buffer and incubated for 15 minutes protected from light. Cells were centrifuged and washed in 1-5 mL MACS buffer 3x. After the final wash/centrifugation, the cell pellet was resuspended in MACS buffer (volume adjusted for cell count) and passed through a 35 μm filter lid (Corning, 352235) into a FACS tube for sorting.

Using single-stain controls from experimental animals and wild-type littermates (TdTomato^-^) for compensation and adjusting gating to remove debris/doublets, the live/CD31^-^/CD45^-^/CD326^+^(EpCAM^+^)/TdTomato^+^ (AEP) population was sorted into a tube containing ‘spiked’ SAGM organoid medium (see ‘*Organoid Medium*’ section below) at 4°C, using a BD FACSAria Fusion cell sorter with a 100 μm nozzle. Yield is approximately 10^5^ AEPs per mouse using this protocol.

### Organoid Medium

To generate ‘spiked’ SAGM medium for mouse lung alveolar organoids, SABM Small Airway Epithelial Cell Growth Basal Medium (Lonza, CC-3119) was combined with the following additives: SAGM Small Airway Epithelial Cell Growth Medium SingleQuots Supplements and Growth Factors (using <u>only</u> the BPE [2 mL], Insulin [0.5 mL], Retinoic Acid [0.5 mL], Transferrin [0.5 mL], and hEGF [0.5 mL] aliquots) (Lonza, CC-4124), Heat Inactivated Fetal Bovine Serum (Corning, 35-011-CV, final concentration 5%), Antibiotic-Antimycotic (Gibco, 15240-062, final concentration 1x), Cholera Toxin from *Vibrio cholerae* (Sigma, C8052, final concentration 25 ng/mL).

### Standard Organoid Plating/Maintenance

AEPs (live/CD31^-^/CD45^-^/CD326^+^[EpCAM^+^]/TdT^+^ cells) sorted from Axin2^creERT2-tDT^ mice were counted using a hemocytometer and resuspended in ‘spiked’ SAGM at a concentration 500 cells/μL. Fibroblasts were prepared (as described above), counted, and resuspended in ‘spiked’ SAGM at a concentration of 5000 cells/μL. For the remaining steps, it was extremely important that all reagents are kept cold/on ice and that bubbles were not introduced to mixtures when pipetting. It is recommended to prepare the plate (Falcon 24 well companion plates [Corning, 353504]), insert Transwells (Falcon Transwell Insert/Permeable Support with 0.4 μm membrane [Corning, 353095]), and place them on ice before use.

For each well of organoids to be plated, 10 μL AEPs (5000 total cells), 10 μL fibroblasts (50000 total cells), and 25 μL ‘spiked’ SAGM were combined (create one master mix of cells and medium for all wells before adding Matrigel) and placed on ice. Corning Matrigel GFR Membrane Matrix (Corning, 356231) was added to the cell mixture (45 μL per well, 1:1 ratio of SAGM to Matrigel) and carefully mixed, then placed back on ice.

For plating, 90 μL of the combined cell/Matrigel mixture was pipetted carefully directly into the center of the Transwell (placed in the companion plate) without introducing bubbles. Organoid plates were incubated at 37°C for 15 minutes, then 500 μL of ‘spiked’ SAGM supplemented with ROCK Inhibitor/Y-27632 Dihydrochloride (Sigma, Y0503, final concentration 0.01 mM) was added beneath the Transwell insert. After 48 hours (and for subsequent media changes), media was replaced every 2 days with ‘spiked’ SAGM <u>without</u> ROCK inhibitor and plates were maintained at 5% CO2 and 37°C.

### AAV6.2FF-Cre Organoids Plating/Maintenance

AAV6.2FF-Cre (titer of 2.779×10^10^ viral genomes [vg]/μL) was generated and characterized *in vivo* as previously described^69,70^. Working dilutions (2.779×10^9^ vg/μL, 2.779×10^8^ vg/μL, and 2.779×10^7^ vg/μL) were generated via serial dilution of viral stocks in ‘spiked’ SAGM and frozen -80°C in single use aliquots.

AEPs (live/CD31^-^/CD45^-^/CD326^+^[EpCAM^+^]/TdT^+^cells) sorted from Axin2^creERT2-tDT^; R26R^EYFP^ mice were counted using a hemocytometer and resuspended in ‘spiked’ SAGM at a concentration 1000 cells/μL. The total cells needed for the desired number of wells were transferred to a new 1.5 mL tube (i.e., 10 wells → 50000 cells → 50 μL cells [1000 cells/μL]). Total cell number per tube, desired MOI (i.e., 1000, 10000, 20000), and known viral titers were used to calculate the volume of virus needed from viral stocks. For each MOI, the calculated volume of virus was added to each cell mixture, mixed, and incubated on ice for 60 minutes. Following viral incubation, ‘spiked’ SAGM and fibroblasts (5000 cells/μL) were added to create a mixture with the same proportions of cells as described above for standard plating of organoids (i.e., for each well – 5000 AEPs + 50000 fibroblasts in 45 μL ‘spiked’ SAGM). The cell mixture was mixed with Matrigel (45 μL/well) and plated/maintained as described above for standard organoids.

#### Organoid Plating with Fibroblasts on Basolateral Side of Transwell

One day prior to organoid plating, fibroblasts were prepared (as described above), counted, and resuspended at a concentration of 50000 cells in 100 μL in fibroblast medium. Transwells were placed in the wells of the companion plate, then the plate was flipped so the Transwells rested on the inside of the lid. The companion plate was removed exposing the basolateral side of the Transwells/filters. 100 μL of the resuspended fibroblast mixture was added to the basolateral side of each Transwell filter. The plate base was placed back on top of the Transwells and was incubated (basolateral side up) at 37°C and 5% CO2 for 4 hours. After incubation, the 100 μL of medium was removed via gentle pipetting (without disturbing the filter) and the plate was flipped to the standard orientation. The Transwells were washed with 500 μL of DPBS (beneath the Transwell insert) then moved to a fresh well/plate with 500 μL fibroblast medium beneath the Transwell insert. After standard isolation of epithelial cells (AEPs) for organoid plating, the Transwells were again washed with 500 μL of DPBS (beneath the Transwell insert), then the epithelial cell/Matrigel mixture (5000 AEPs in 45 μL ‘spiked’ SAGM + 45 μL Matrigel) was added to the apical side of the Transwell filter. Organoids were maintained as described above for standard organoids.

### Fixation/Processing for Sections/Histology/H&E

Organoids were washed with 500 μL PBS, above and below Transwells. After removing PBS, 500 μL of 4% PFA was added above and below the Transwell for fixation overnight at 4°C. Transwells were washed 5x (above and below) with PBS. Using a small knife or scalpel, the Transwell filter and Matrigel plug/organoids were cut out of the Transwell and placed on parafilm. Using forceps, the Transwell filter was carefully removed from the Matrigel plug/organoids [note: older organoid cultures are more likely to adhere to the filter]. Using a transfer pipet, HistoGel (Epredia, HG4000012) (pre-heated to a liquid consistency) was added on top of the Matrigel plug/organoids until covered on all sides. Once solidified (∼15-30 minutes), the sample was transferred to a tissue processing cassette (Fisher, 15-182-702A). Once in cassettes, samples were processed as described for whole-lung processing above for paraffin embedding and sectioning, H&E, and immunofluorescence of paraffin sections.

### Isolation of Organoids for Whole-Mount Immunofluorescence

Previously established whole-mount organoid staining protocols from Dekkers, et al.^28^ were adapted for mouse lung alveolar organoids using the following modifications. All steps used cut or wide-bore pipette tips. All steps after first wash and prior to fixation were performed on ice/with chilled reagents and used cut/wide bore pipette tips coated in 1% BSA in PBS.

Briefly, Transwells were washed (above and below) with 500 μL of room temperature PBS. 500 μL of Cell Recovery Solution (Corning, 354253) was added to each Transwell, and a cut pipette tip was used to mechanically disrupt the Matrigel – the mixture was pipetted up and down and transferred to a new 24 well plate. Each Transwell was washed with an additional 250 μL of cell recovery solution and added to the new plate. The plate was incubated on ice (gel ice packs were optimal) on an orbital/horizontal shaker for 60 minutes. The organoid mixture was transferred to a 15 mL conical pre-coated in 1% PBS-BSA. Each well was washed with 500 μL of 1% PBS-BSA and added to the 15 mL conical. Wells from the same experimental condition (up to 4 wells) were combined in one conical. Conicals were filled to 10 mL with ice cold PBS and centrifuged at 70g for 5 minutes at 4°C. The supernatant was removed very carefully [note: if the organoid pellet is not compact/tight, the entire pellet may be lost with suction due to loose matrix. If Matrigel was still visible, the organoid pellet was gently resuspended in 1 mL of ice cold 1% PBS-BSA and centrifuged again at 70g for 5 min at 4 °C. After careful removal of the supernatant, the organoid pellet was resuspended in 1 mL of 4% PFA and incubated at 4°C for 45 minutes (resuspending once halfway through incubation). For permeabilization, conicals were filled to 10 mL with 0.1% PBS-Tween and incubated overnight at 4°C (alternate permeabilization option: for Click-iT protocols or shorter permeabilization, remove PFA and incubate in 0.25% Triton X-100 for 20 minutes at room temperature).

### Whole-Mount Blocking and Immunofluorescence

After isolation, fixation, and permeabilization, organoids were centrifuged at 70g for 5 min at 4°C, resuspended in 500 μL of 5% Normal Donkey Serum (Jackson ImmunoResearch, 017-000-121) in 0.1% PBS-Triton X-100, and transferred to a 24-well plate for blocking. Organoids were incubated at room temperature on an orbital shaker for 1-2 hours.

After blocking, the supernatant was removed from each well without disturbing organoids [note: supernatant was removed more easily when plate was placed at a 45° angle for 5-10 minutes, to allow organoids to settle to bottom edge of well]. Primary antibodies (see supplemental table 1) were diluted to a final concentration of 1:100 in 5% Normal Donkey Serum in 0.1% PBS-Triton X-100 (approximately 200-250 μL total) and incubated overnight at 4°C on an orbital shaker. For this protocol, a ‘quick wash’ was defined as adding 1 mL of organoid wash buffer (0.2% BSA, 0.1% Triton X-100 in PBS)^28^ and immediately allowing organoids to settle/removing the wash, and a ‘long wash’ was defined as adding 1 mL of organoid wash buffer placing the plate on an orbital shaker for 1-2 hours before allowing organoids to settle/removing the wash. After primary antibody staining, one ‘quick wash’ and three ‘long washes’ were performed. Then, secondary antibodies (see supplemental table 1) were diluted to a final concentration of 1:200 in 5% Normal Donkey Serum in 0.1% PBS-Triton X-100 (approximately 200-250 μL total) and incubated overnight at 4°C on an orbital shaker. For this step and all subsequent steps, samples were covered/protected from light to prevent photobleaching. After secondary antibody staining, one ‘quick wash’ was performed, then organoids were incubated in DAPI (Invitrogen, D1306, final concentration of 1:1000) in 5% Normal Donkey Serum in 0.1% PBS-Triton X-100 (approximately 200-250 μL total) for 15 minutes. After removing supernatant, one ‘quick wash’ and three ‘long washes’ were performed.

### Whole-Mount Clearing and Mounting

Following the final wash after immunostaining, as much wash buffer as possible was removed from each well and organoids were transferred to a 1.5 mL tube. Organoids were centrifuged at 70g for 5 min at 4 °C and as much supernatant as possible was removed without disturbing the organoids. Using a cut or wide-bore pipette tip, organoids were gently resuspended in room temperature fructose-glycerol clearing solution (60% vol/vol glycerol + 2.5 M fructose)^28^. Depending on organoid volume, ∼50-200 μL of clearing solution was used. Organoids were left to clear for at least 1 day (and as long as several months) at 4°C before mounting.

Prior to preparing slides, cleared organoids were allowed to equilibrate to room temperature. Organoids were mounted as described previously^28^– briefly, two pieces of double-sided tape were applied to a microscope slide approximately 25-30 mm apart, perpendicular to the length of the slide (for larger organoids, additional layers of tape can be used). Using a PAP pen (Abcam, ab2601), a square was drawn between the two pieces of tape. Using a cut P200 pipette tip, approximately 20 μL of organoids in clearing solution was placed in the middle of the drawn square, avoiding bubbles. A #1.5 Gold Seal 3419 Cover Glass (Electron Microscopy Sciences, 63790-01), was applied over the organoids, bridging the two pieces of tape. Slides were imaged immediately or stored at 4°C.

### Whole-Mount Click-iT EdU Staining

For whole-mount Click-iT EdU staining, the standardized commercial protocol for Click-iT EdU Cell Proliferation Kit for Imaging, Alexa Fluor 488 dye (Invitrogen, C10337) was combined with our optimized whole-mount immunofluorescence protocol. Kit reagents were prepared as directed in commercial protocols. Briefly, 48 hours prior to harvest/fixation, organoid medium was replaced with ‘spiked’ SAGM supplemented with EdU (final concentration of 10 μM) from the commercial kit. After EdU incubation, all steps described in ‘*Isolation of Organoids for Whole-Mount Immunofluorescence*’ were performed. Next, steps for EdU detection from the commercial kit’s standardized protocol with kit reagents were followed (i.e., 30-minute incubation of “Click-iT Reaction Cocktail” at room temperature, followed by 1 mL wash with 3% PBS-BSA). Following EdU detection, whole-mount immunofluorescence was performed as described in the ‘*Whole-Mount Blocking and Immunofluorescence’* and ‘*Whole-Mount Clearing and Mounting*’ sections above.

### Whole-Mount Click-iT TUNEL Staining

For whole-mount Click-iT TUNEL staining, the standardized commercial protocol for Click-iT Plus TUNEL Assay for In Situ Apoptosis Detection, Alexa Fluor 488 dye (Invitrogen, C10617) was combined with our optimized whole-mount immunofluorescence protocol. Kit reagents were prepared as directed in commercial protocols. All steps described in ‘*Isolation of Organoids for Whole-Mount Immunofluorescence*’ were performed. Organoids were washed twice with DI H2O. Next, steps for ‘TdT Reaction’ and ‘Click-iT Plus Reaction’ from the commercial kit’s standardized protocol with kit reagents were followed (i.e., 60-minute incubation of “TdT Reaction Mixture” at 37°C, followed by 2 washes with 3% PBS-BSA, and a 30-minute incubation of “Click-iT Plus TUNEL reaction cocktail” at 37°C). Organoids were washed twice with 3% PBS-BSA, then whole-mount immunofluorescence was performed as described in the ‘*Whole-Mount Blocking and Immunofluorescence’* and ‘*Whole-Mount Clearing and Mounting*’ sections above.

### Hoechst and Live Imaging Preparation

For live imaging of organoids grown with PDGFRα^EGFP^ fibroblasts, Hoechst 33342 (Invitrogen, H3570) was diluted 1:10000 in ‘spiked’ SAGM and 500 μL was added above and below the Transwell and incubated at 37°C for 30-45 minutes. Using a small knife or scalpel, the Transwell filters and Matrigel plug/organoids were cut out of the Transwells and placed into a coverslip bottom dish (MatTek, P35G-1.5-20-C). For some samples, the entire Matrigel plug/filter was imaged, and for others the Matrigel plug and filter were separated and imaged independently. Samples were covered in ‘spiked’ SAGM and cover slipped (MatTek, PCS-1.5-18) prior to imaging on an inverted confocal microscope.

### Imaging

Brightfield H&E images were acquired on a Nikon Eclipse NiE Upright Widefield Microscope (Nikon DS-Fi3 Camera – with a Plan Apo VC 20x DIC N2 objective). Fluorescent images were acquired on Nikon A1 inverted LUNV and Nikon A1R inverted LUNV confocal microscopes using the following objectives: Plan Apo λ 10x, Plan Apo λ 20x, Apo LWD 20x WI λS (water immersion), Apo LWD 40x WI λS DIC N2 (water immersion), and SR HP Plan Apo λ S 100xC Sil (silicone immersion). Second harmonic generation was performed to visualize fibrillar collagens I and II using a Nikon FN1 Upright Multiphoton microscope using the following objectives: Plan Apo VC 20x DIC N2 and Apo LWD 25x 1.10W DIC N2. Images were processed in Nikon Elements with minimal, global adjustment of LUTs for acquired channels.

### Organoid Plate Imaging/Cytation Imager

For whole well imaging, plates were loaded into a Cytation 5 Imager (BioTek, CYT5PV) configured with a CO2 gas controller (BioTek, 1210012). Plates were maintained at 5% CO2 and 37°C during imaging using *Cytation Gen5 Microplate Reader and Imager Software* (BioTek, version 3.08.01). Protocols specific to Falcon 24 well companion plates (Corning, 353504) and Falcon Transwell Insert/Permeable Support with 0.4 μm membrane (Corning, 353095) were established and used to take brightfield and fluorescent (GFP) 4x tile scans at 10 z-steps (∼50 μm per step). 4x tile scans were used to generate z-projections. Individual tile scans and z-projections were used for further quantification.

### Organoid Quantification

Z-projections of stitched 4x images from each well were loaded into a custom FIJI-macro (run in FIJI/ImageJ v1.53) to count organoids per well, GFP^+^ organoids per well, and organoid area. This macro allowed for batch analysis of each experiment, reducing subjectivity of counts. Briefly, given specific input parameters, the macro contained commands to: set the scale based on the diameter of each Transwell, subtract background, adjust image threshold, convert to mask, analyze particles/count objects meeting a specific threshold, and export data. Data was imported into GraphPad Prism 9.0 for analysis. T tests were used for comparison of 2 groups, and ANOVA with prespecified multiple comparisons was used to compare 3 or more groups.

### Electron Microscopy

Fixation, sectioning, and acquisition of electron micrographs of alveolar cells was performed as previously described^89^.

### Organoid Dissociation and Preparation of Single Cell Suspension for scRNAseq and scATACseq

Transwells were washed (above and below) with 1 mL of PBS. 60 μL of organoid digest buffer (Dispase [Corning, 354235, undiluted, 50 U/mL], DNase I [GoldBio, D-301, final concentration 0.25mg/mL], Collagenase Type I [Gibco, 17100017, final concentration 4800 U/mL)]) was added and Matrigel plugs were gently disrupted and pipetted using a cut or wide-bore pipette tip. Organoids were incubated in digest buffer for 30 minutes at 37°C. Following incubation, the digested organoid mixture was pipetted several times and transferred to a low-binding 1.5 mL tube (3 wells of same experimental condition combined into each tube). 500 μL of cold PBS was added to each tube and incubated on ice for 5-10 minutes. Samples were centrifuged at 500g for 5 minutes at 4°C. Supernatant was removed carefully and sample was washed with 1 mL of cold DPBS (Gibco, 14190-094). Following centrifugation at 500g for 5 minutes at 4°C and removal of supernatant, samples were resuspended in 60 μL of 0.25% Trypsin-EDTA (Gibco, 25200-056) and incubated for 30 minutes at 37°C. 1 mL of ice-cold PBS was added to each tube, samples were centrifuged at 500g for 5 minutes at 4°C, and the supernatant was carefully removed. Samples were washed 2x in 1 mL of cold 0.04% PBS-BSA and centrifuged at 500g for 5 minutes at 4°C. Following removal of the supernatant, the samples were resuspended in 100 μL of 0.04% PBS-BSA. Prior to filtering cells, 40 μm Flowmi Cell Strainers (Bel-Art, H13680-0040) were equilibrated by passing 100 μL of 0.04% PBS-BSA through the strainer using a P1000 pipette tip. The 100 μL cell suspension was then pipetted through the 40 μm Flowmi Cell Strainer. Cells were counted manually using a hemocytometer and resuspended at a concentration of 1000 cells/μL prior to processing for scRNAseq.

### Nuclei Isolation from Organoids for scATACseq

Using the same filtered cell suspension generated for scRNAseq, the standard 10x Genomics protocol for ‘Nuclei Isolation for Single Cell ATAC Sequencing’ (CG000212 Revision B) was followed. Briefly, the single cell suspension was centrifuged at 500g for 5 minutes at 4°C. Following removal of the supernatant, 100 μL of ATAC lysis buffer (from standard 10x Genomics protocol, CG000212 Revision B) was added, gently mixed, and incubated on ice for 4-4.5 minutes. Immediately following incubation, 1 mL of chilled ATAC wash buffer (from standard 10x Genomics protocol) was added and gently mixed. Nuclei were centrifuged at 500g for 5 minutes at 4°C, supernatant was removed, and nuclei were resuspended in 100 μL of 1x nuclei buffer (diluted from 20x nuclei buffer [10x Genomics, 2000153/2000207]). Prior to filtering nuclei, 40 μm Flowmi Cell Strainers were equilibrated by passing 100 μL of nuclei buffer through the strainer using a P1000 pipette tip. The 100 μL of nuclei suspension was then pipetted through the 40 μm Flowmi Cell Strainer. Nuclei were counted manually using a hemocytometer and resuspended at a concentration of 5000 nuclei/μL prior to processing for scATACseq.

### Sequencing/Library Preparation

From each single cell or single nuclear preparation described above, a maximum of 16,000 cells or nuclei were loaded into on channel of a 10x Genomics Chromium system by the Cincinnati Children’s Hospital Medical Center Single Cell Sequencing Core. Libraries for RNA (v3) and ATACseq (v2) were generated following the manufacturer’s protocol. Sequencing was performed by the Cincinnati Children’s Hospital DNA Sequencing Core using Illumina reagents. Raw Sequencing data was aligned to the mouse reference genome mm10 with CellRanger 3.0.2 to generate expression count matrix files. To detect YFP expressing cells following Cre-mediated activation, a YFP contig was added to the mm10 genome following 10x Genomics “Build a Custom Reference” instructions(https://support.10xgenomics.com/single-cell-gene-expression/software/pipelines/latest/using/tutorial_mr) with modifications. Briefly, a custom EYFP .fasta file was generated using the ‘EYFP’ segment (682-1389) of the pEYFP-N1 plasmid sequence available through Addgene. This sequence was integrated into the standard mm10 assembly available from Ensembl to create a reference compatible for alignment with the CellRanger pipeline described above.

### scRNAseq Analysis and Visualization

For RNAseq analysis, output data from CellRanger was partitioned into spliced and unspliced reads using Velocyto^90^. Velocyto output files were loaded into Seurat 4.0 using SeuratWrappers and SeuratDisk using the *ReadVelocity* command and spliced transcripts were used as the expression input to *SCTransform*. Cells with less than 2000 or more than 8000 features were filtered and cells were clustered using the standard Seurat workflow. Putative doublets were identified and removed using DoubletFinder^91^, and libraries from individual time points and treatments were integrated using *SelectIntegrationFeatures* and *IntegrateData* commands in Seurat. Following integration, cells were re-clustered, UMAP project generated, and samples identified based on expression similarity to published data as described in the Results. Module scoring was performed using *AddModuleScore* function in Seurat for gene sets indicated in the figures. For lineage inference, these Seurat objects were directly used for Slingshot^42^ pseudotime inference, and were converted to a h5ad file using the *SaveH5Seurat* command. These h5ad file were used as input to scVelo^43^ and CellRank^92^ in Python 3.9.12 in Spyder following the standard pipeline (scVelo.readthedocs.io) to generate RNA velocity mapped to the Seurat UMAP and cell populations. For ligand-receptor analysis, we used CellChat^35^ (https://github.com/sqjin/CellChat) v1.1 using the *SecretedSignaling* subset of the Mouse CellChatDB, with default parameters. Visualizations were generated with these tools and ggplot2.

#### scATACseq Analysis and Visualization

For ATACseq analysis, CellRanger output was loaded into ArchR^45^ and Arrow files were generated per package defaults. Clusters were generated based on ATACseq parameters and named based on evaluation of integrated gene expression from the paired Seurat RNA object. Peak calls for regions of open chromatin were generated from pseudobulk analysis of each cell state followed by peak calling in MACS2. Differential open chromatin peaks were identified based on FDR <0.01 and Log2FC >=1 between cell states. Visualizations were generated using standard ArchR commands.

#### Transcriptional Regulatory Networks and Visualization

For TRN inference, scRNAseq gene expression data and scATACseq chromatin accessibility data from each epithelial cell population was used as input for TRN analysis as previously reported^24^. Enriched transcriptional regulators were identified per cell type by performing Fisher’s Exact Test to compare observed vs expected number of genes regulated in a cell type based on the TRN model, and visualizations were generated in Illustrator with details as noted in the Figure Legends.

---

**Supplemental Table 1a.** Antibodies for Organoid Whole-Mount Immunofluorescence:

| Primary Antibody (1:100 – unless otherwise specified) | Secondary Antibody (1:200) |
| --- | --- |
| anti-Sftpc (Guinea Pig, Seven Hills Bioreagents, GP992) | Goat anti-Guinea Pig IgG Alexa Fluor 555 (Invitrogen, A21435)<br>Goat anti-Guinea Pig IgG Alexa Fluor 647 (Invitrogen, A21450) |
| anti-RAGE (Rat, R&D Systems, MAB1179) | Donkey anti-Rat IgG Alexa Fluor 488 (Invitrogen, A21208)<br>Donkey anti-Rat IgG Alexa Fluor 568 (Abcam, ab175708) |
| anti-TTF1/Nkx2-1 (Rabbit, Seven Hills Bioreagents, Rb1231) | Donkey anti-Rabbit IgG Alexa Fluor 647 (Invitrogen, A31573) |
| anti-Ki67 (Rabbit, Abcam, ab16667 [SP6]) | Donkey anti-Rabbit IgG Alexa Fluor 647 (Invitrogen, A31573) |
| anti-GFP (Chicken, Abcam, ab13970) | Goat anti-Chicken IgG Alexa Fluor 488 (Invitrogen, A11039) |
| anti-CD45 (Rat, BioLegend, 103103) | Rabbit anti-Rat IgG Alexa Fluor 488 (Invitrogen, A21210) |
| anti-E-Cadherin (Rat, R&D Systems, MAB7481 – 1:250) | Donkey anti-Rat IgG Alexa Fluor 568 (Abcam, ab175708) |
| anti-Krt8 (Rat, DSHB, TROMA-I – 1:50) | Donkey anti-Rat IgG Alexa Fluor 568 (Abcam, ab175708) |

**Supplemental Table 1b.**
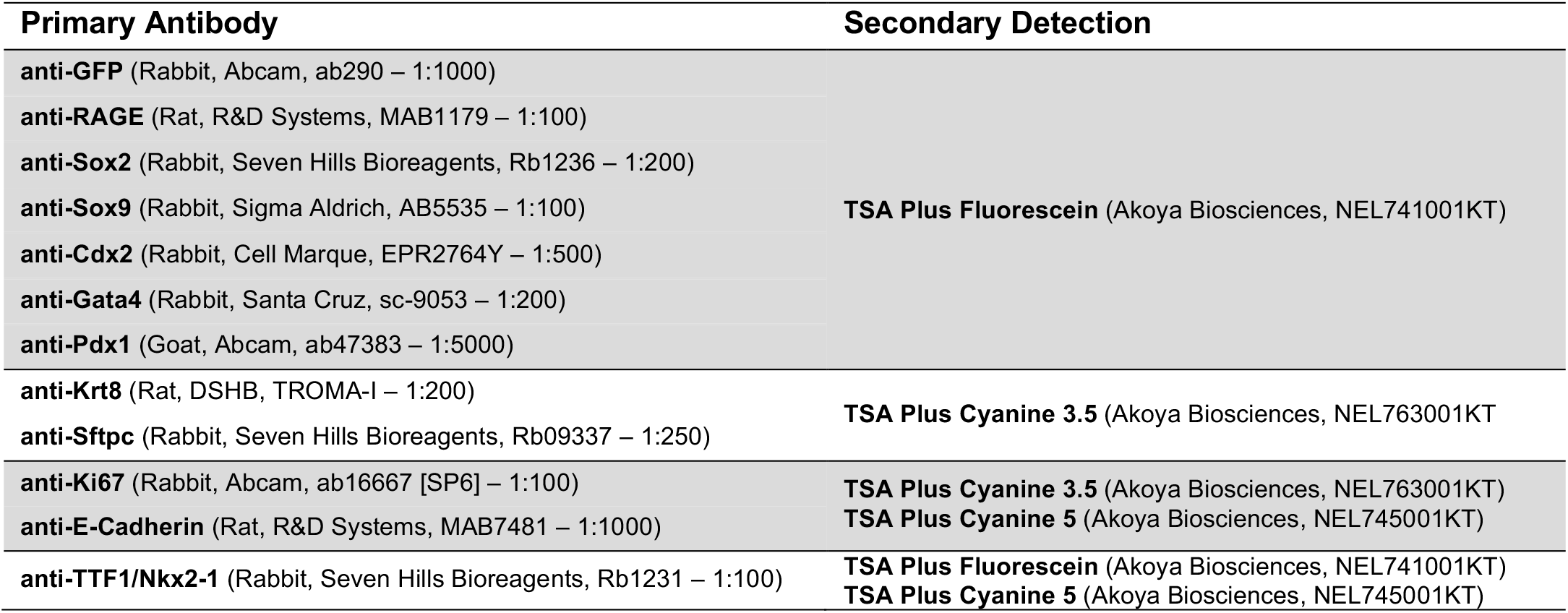
Antibodies for Immunofluorescence on Paraffin Sections:

**Supplemental Table 2.** scRNAseq Output Summary:

| Sample ID | Day 14 | Day 21 | Day 28 | Day 28 AAV Control | Day 28 Nkx2-1 KO |
| --- | --- | --- | --- | --- | --- |
| Estimated Number of Cells | 9,908 | 9,519 | 5,546 | 9,184 | 11,057 |
| Mean Reads per Cell | 40,509 | 42,230 | 69,615 | 63,819 | 44,557 |
| Median Genes per Cell | 3,606 | 3,637 | 4,431 | 2,766 | 2,925 |
| Number of Reads | 401,364,390 | 401,993,978 | 386,086,102 | 586,113,691 | 492,668,484 |
| Valid Barcodes | 97.9% | 98.2% | 98.2% | 97.60% | 97.70% |
| Sequencing Saturation | 23.4% | 28.9% | 41.9% | 28.60% | 28.10% |
| Q30 Bases in Barcode | 95.8% | 95.8% | 95.8% | 96.70% | 96.70% |
| Q30 Bases in RNA Read | 93.3% | 93.6% | 93.7% | 94.20% | 93.90% |
| Q30 Bases in UMI | 95.4% | 95.4% | 95.4% | 96.20% | 96.20% |
| Reads Mapped to Genome | 90.9% | 91.9% | 93.0% | 82.80% | 85.60% |
| Reads Mapped Confidently to Genome | 86.7% | 88.0% | 89.2% | 78.30% | 81.50% |
| Reads Mapped Confidently to Intergenic Regions | 6.0% | 6.1% | 7.2% | 8.50% | 12.20% |
| Reads Mapped Confidently to Intronic Regions | 12.7% | 11.1% | 10.4% | 11.80% | 10.40% |
| Reads Mapped Confidently to Exonic Regions | 67.9% | 70.7% | 71.6% | 58.00% | 58.90% |
| Reads Mapped Confidently to Transcriptome | 64.5% | 67.7% | 68.8% | 55.20% | 56.40% |
| Reads Mapped Antisense to Gene | 2.0% | 1.6% | 1.4% | 1.90% | 1.20% |
| Fraction Reads in Cells | 76.7% | 78.2% | 83.7% | 48.90% | 84.40% |
| Total Genes Detected | 20,300 | 20,134 | 19,308 | 22,830 | 17,723 |
| Median UMI Counts per Cell | 13,384 | 14,538 | 22,287 | 9,140 | 10,571 |

**Supplemental Table 3.** scATACseq Output Summary:

| Sample ID | Day 14 | Day 21 | Day 28 |
| --- | --- | --- | --- |
| Annotated Cells | 13,300 | 11,765 | 4,535 |
| bc_q30 Bases Percent | 89.4% | 90.0% | 78.4% |
| Cellranger-ATAC Version | 1.2.0 | 1.2.0 | 1.2.0 |
| Cells Detected | 13,300 | 11,765 | 4,535 |
| Percent Cut Fragments in Peaks | 47.0% | 34.7% | 60.3% |
| Percent Fragments nfr | 67.8% | 71.0% | 68.6% |
| Percent Fragments nfr or nuc | 94.4% | 96.1% | 93.2% |
| Percent Fragments nuc | 26.6% | 25.1% | 24.6% |
| Percent Fragments Overlapping Peaks | 49.1% | 36.7% | 62.8% |
| Percent Fragments Overlapping Targets | 63.4% | 55.6% | 74.6% |
| Percent Mapped Confidently | 79.3% | 76.9% | 69.3% |
| Percent Waste Chimeric | 0.6% | 0.4% | 0.5% |
| Percent Waste Duplicate | 13.6% | 7.7% | 10.6% |
| Percent Waste lowmapq | 5.2% | 4.0% | 3.4% |
| Percent Waste Mitochondrial | 0.3% | 0.3% | 0.7% |
| Percent Waste No Barcode | 1.7% | 1.7% | 5.7% |
| Percent Waste Non Cell Barcode | 46.6% | 66.9% | 54.0% |
| Percent Waste Overall Nondup | 55.1% | 73.6% | 66.0% |
| Percent Waste Total | 68.7% | 81.3% | 76.6% |
| Percent Waste Unmapped | 0.7% | 0.4% | 1.6% |
| Median Fragments Per Cell | 10,746 | 7,700 | 13,310 |
| Median per Cell Unique Fragments at 30000 RRPC | 4,135.0 | 2,578.2 | 4,064.0 |
| Median per Cell Unique Fragments at 50000 RRPC | 6,501.4 | 4,095.3 | 6,437.5 |
| Num_Fragments | 6.13E+08 | 6.26E+08 | 2.73E+08 |
| r1_q30 Bases Percent | 94.4% | 94.3% | 86.6% |
| r2_q30 Bases Percent | 94.0% | 94.0% | 85.2% |
| si_q30 Bases Percent | 92.4% | 92.5% | 76.4% |
| Total Usable Fragments | 1.92E+08 | 1.17E+08 | 6.40E+07 |
| TSS Enrichment Score | 6.360 | 4.651 | 8.864 |

**Supplemental Figure 1.**
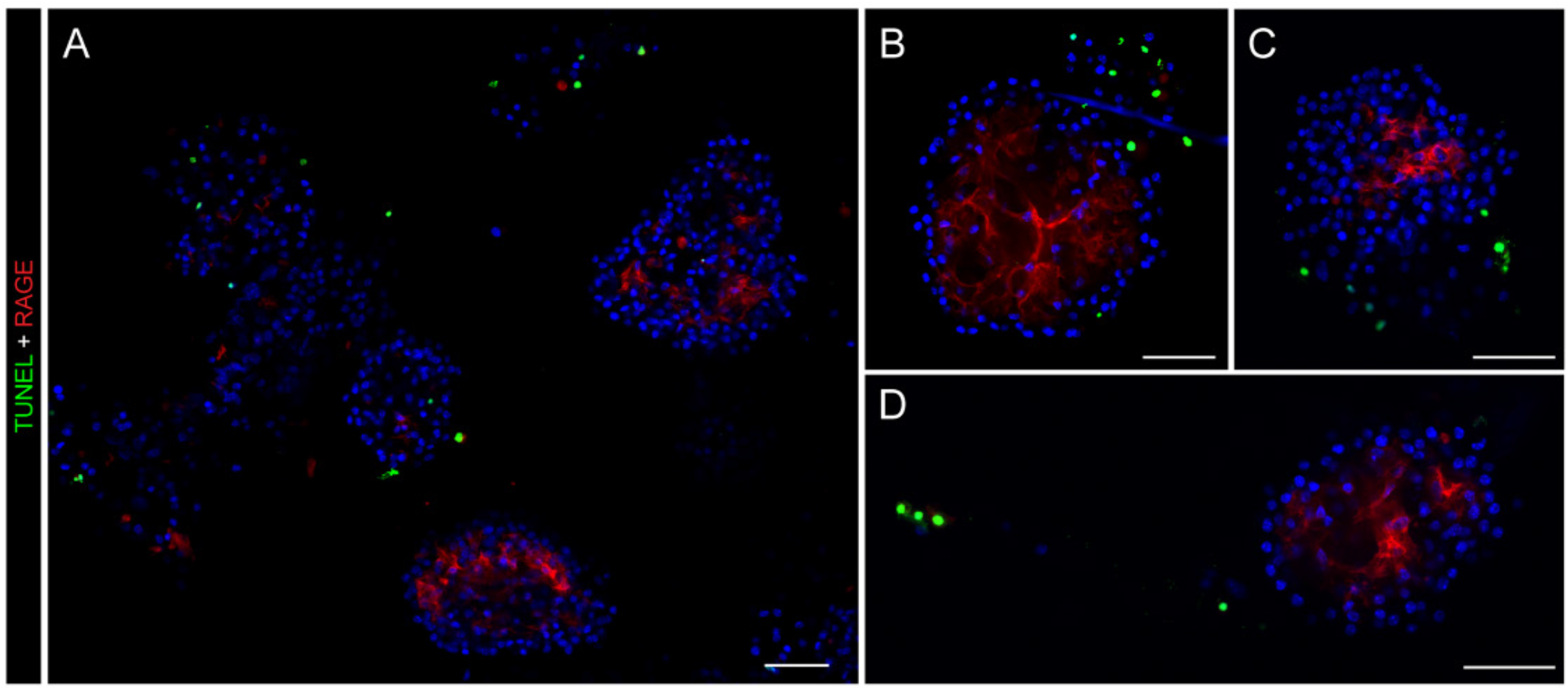
TUNEL^+^ cells largely confined to cell clumps and debris outside organoids. (A-D) Whole-mount immunofluorescence and Click-iT TUNEL staining showing lack of TUNEL+ cells within mature, healthy day 25 organoids. [*Scale bars = 50 μm*]; (*TUNEL = Terminal deoxynucleotidyl transferase dUTP nick end labeling [indicator of DNA strand breaks/apoptosis]; RAGE = Receptor for Advanced Glycation End-products [AT1 cell marker]*)

**Supplemental Figure 2.**
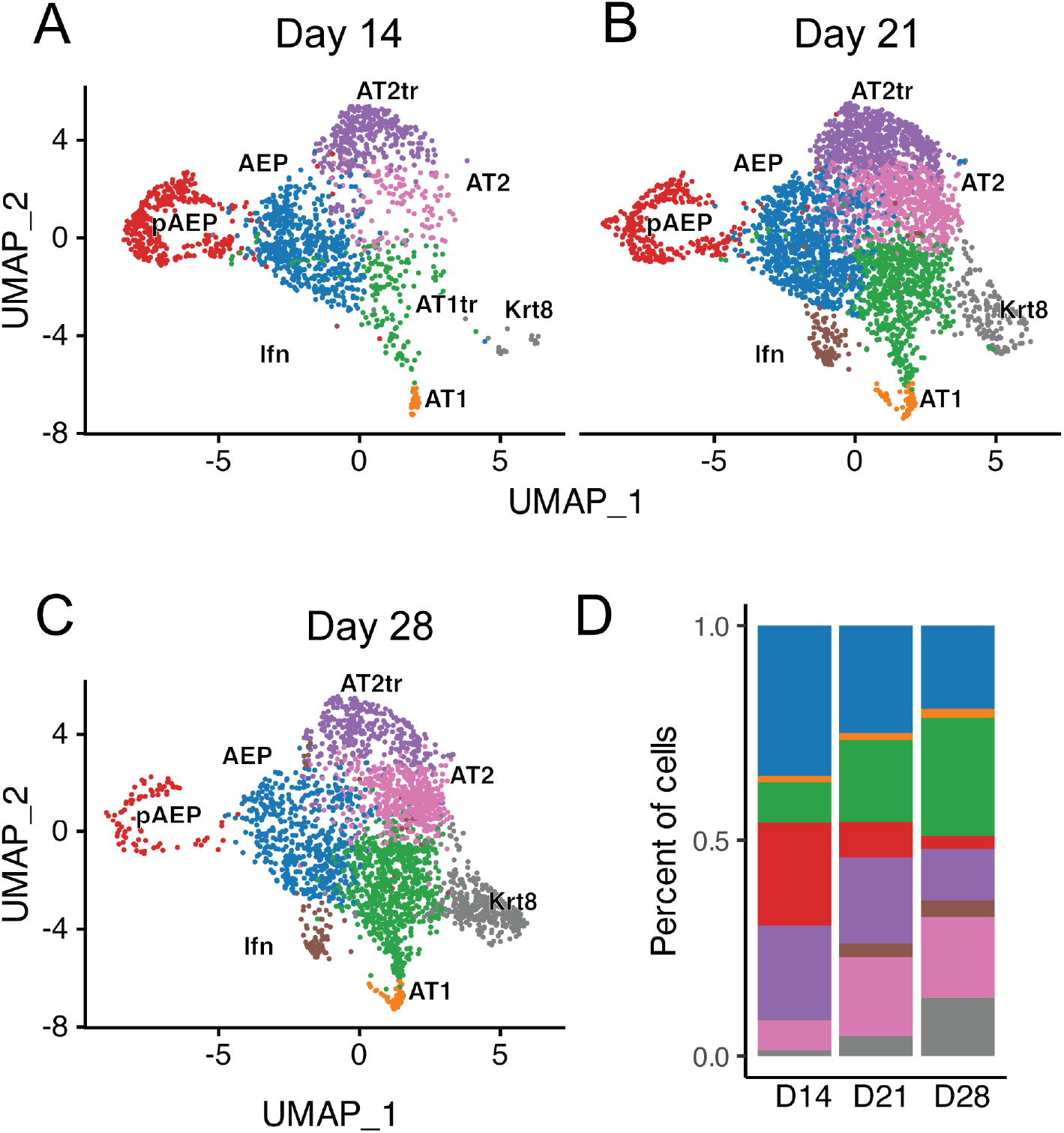
Evolution and contribution of epithelial cell states per timepoint in AEP-O. A-C) UMAP projections (left) demonstrating relative detected cells for each epithelial cell state at day 14 (A), day 21 (B), and 28 (C). D) Quantification of cell population abundance at each time point.

**Supplemental Figure 3.**
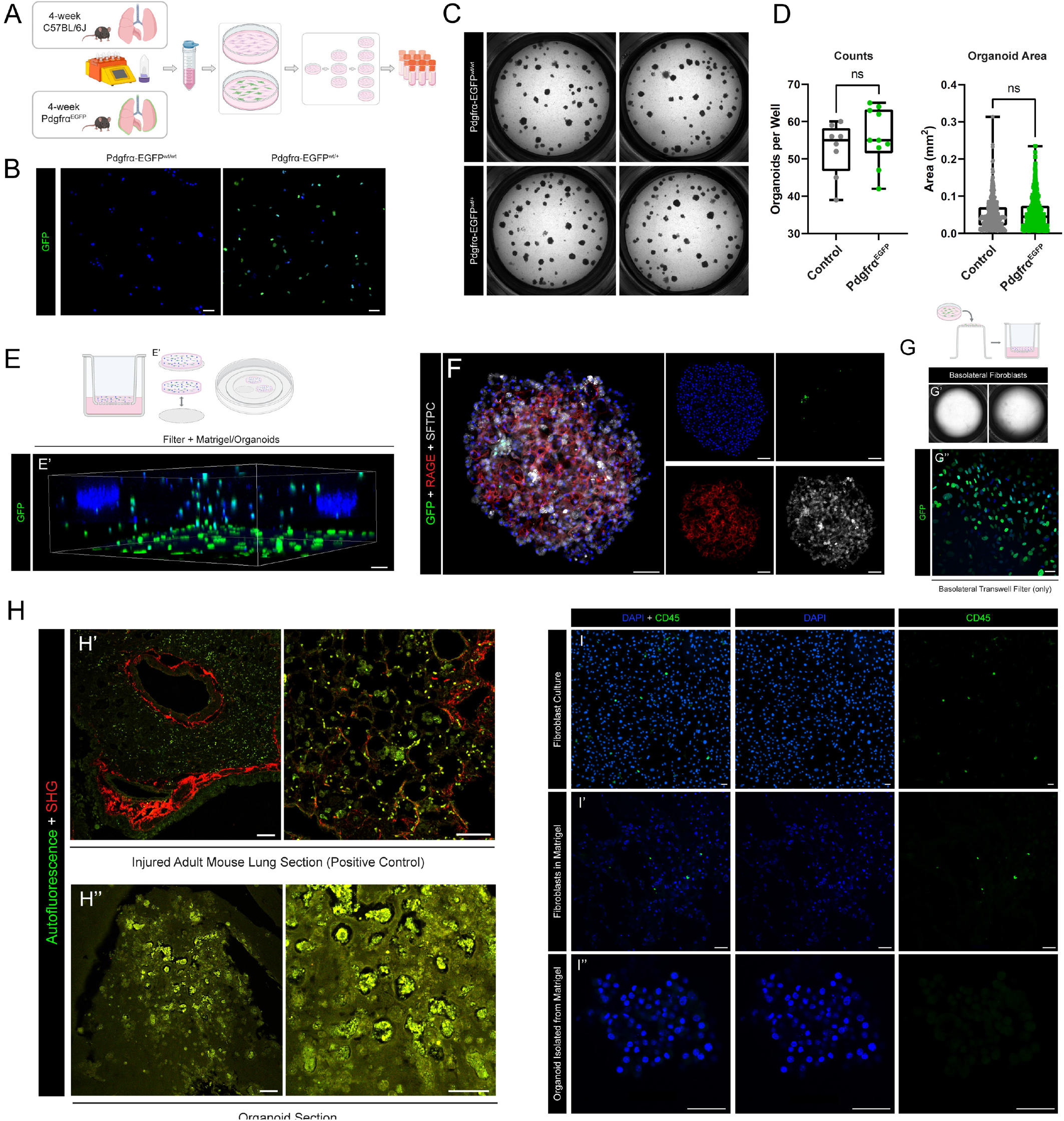
Localization of mesenchymal and immune cells in wells surrounding AEP-O. (A) Generation of fibroblast stocks from control (C57BL/6J) and PDGFRα^EGFP^ mice. (B) Imaging of control and GFP fibroblast stocks at P2 (second passage). (C) Whole-well scans comparing day 35 AEP-derived organoids grown from control and PDGFRα^EGFP^ fibroblasts. (D) Quantification comparing counts and size of day 35 AEP-derived organoids grown from control and PDGFRα^EGFP^ fibroblasts. (E) Schematic of experimental set-up of live imaging; 3D reconstruction of live day 20 organoids and PDGFRα^EGFP^ fibroblasts stained with Hoechst; (E’) 3D reconstruction of confocal z-stacks of Matrigel/organoids with the Transwell filter, showing the majority of GFP^+^ fibroblasts are growing on the filter and not within organoids; (F) Whole-mount immunofluorescence of organoids grown with PDGFRα^EGFP^ fibroblasts showing lack of PDGFRα^+^ cells within day 35 organoids. (G) Cultures with fibroblasts on the basolateral side of the filter and AEPs on the apical side of the filter do not grow organoids (G’ – 31-day culture), although fibroblasts do persist on the basolateral side of the filter after 31 days (G’’). (H) Confocal imaging utilizing second harmonic generation to show lack of fibrillar collagen I and II in paraffin sections of organoids from long term culture (H’’) (injured adult mouse lung section as positive control [H’]). (I-I’’) Immunofluorescence staining showing presence of contaminating immune (CD45^+^) population (green) in fibroblast stocks (I). Whole-mount immunofluorescence of day 28 Matrigel/fibroblast mixture showing presence of contaminating immune (CD45^+^) population (green) outside organoids in Transwell cultures (I’). Whole-mount immunofluorescence of day 28 organoids showing lack of immune (CD45^+^) population (green) inside organoids (I’’). (*ns = p > 0*.*05)* [*Scale bars = 50 μm*].

**Supplemental Figure 4.**
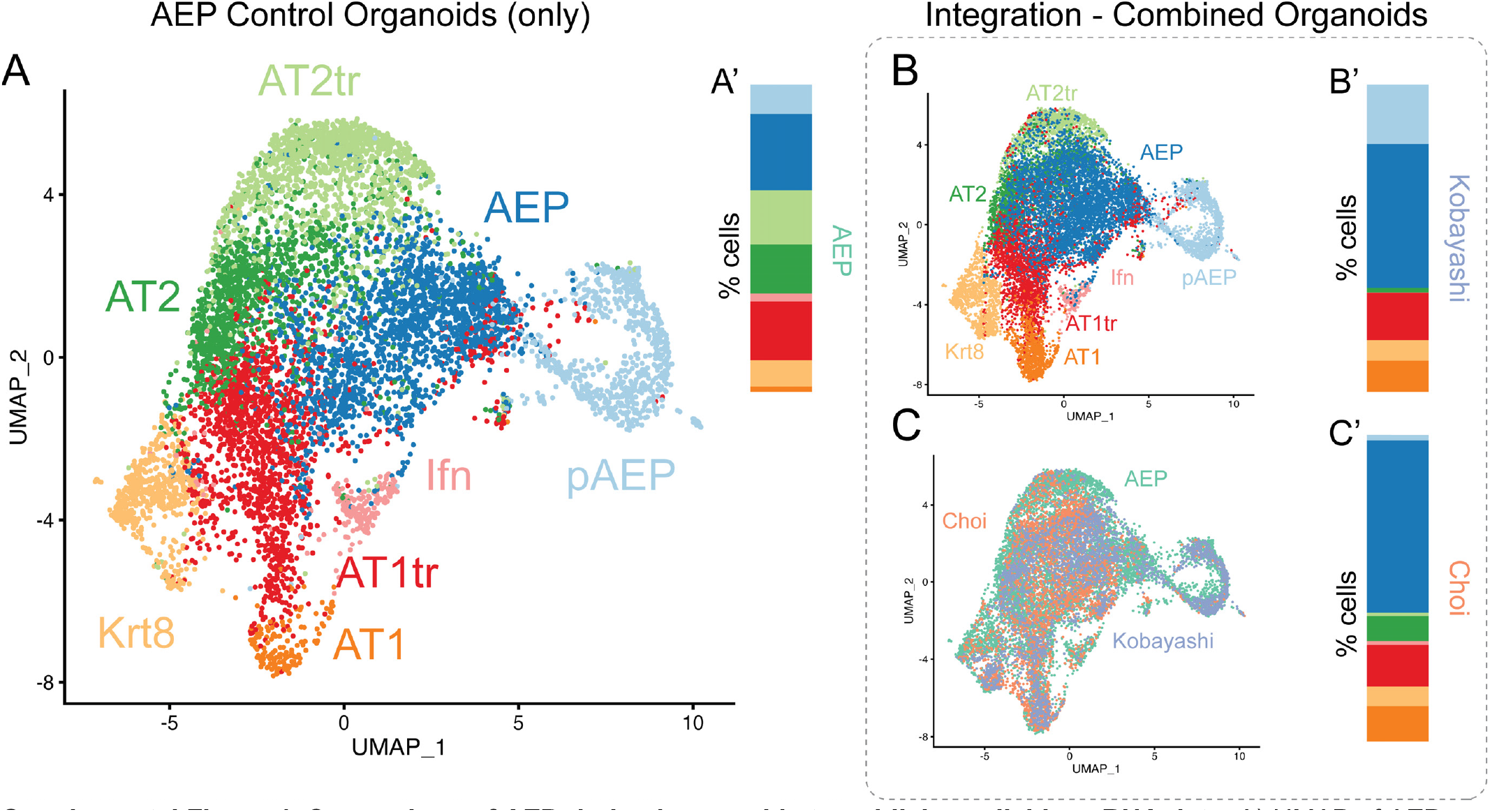
Comparison of AEP-derived organoids to publicly available scRNA data. A) UMAP of AEP organoids used as basis of integration and labels. E) Integrated data from all three organoid datasets, labeled by cell type (B) or dataset of origin (C). Composition of each organoid dataset is shown in A-C’.

**Supplemental Figure 5.**
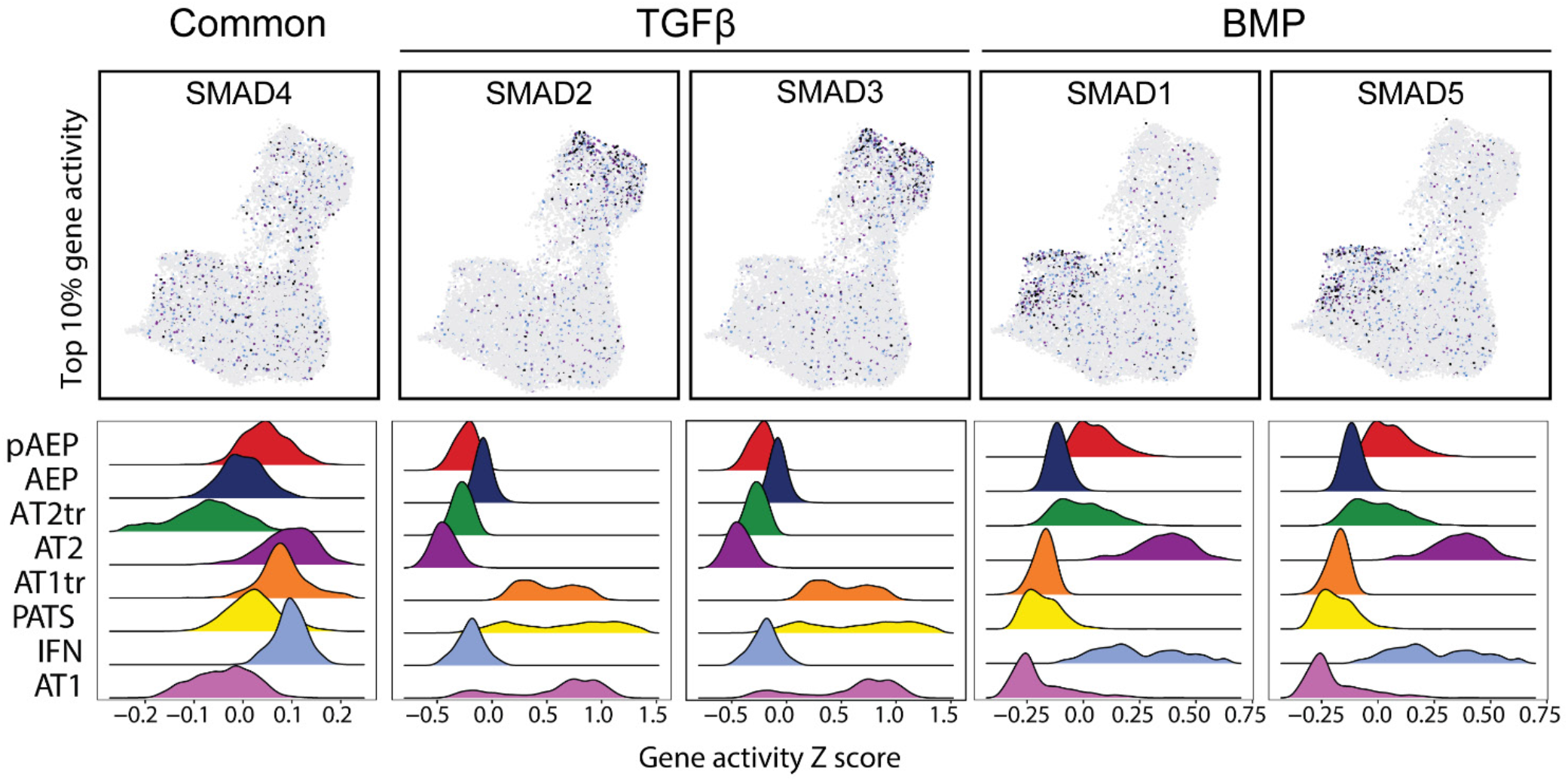
Transition of SMAD-regulated gene expression in AT2 to AT1 transitions in AEP-O. Top row shows gene activity of SMAD target genes overlayed on scATACseq UMAP (compare to Figure 4A). Bottom row shows gene activity Z score of SMADs in each cell population in AEP-O; BMP signaling is predicted to be higher in AT2 cells, while TGFβ signaling predominates in AT1 cells.

**Supplemental Figure 6.**
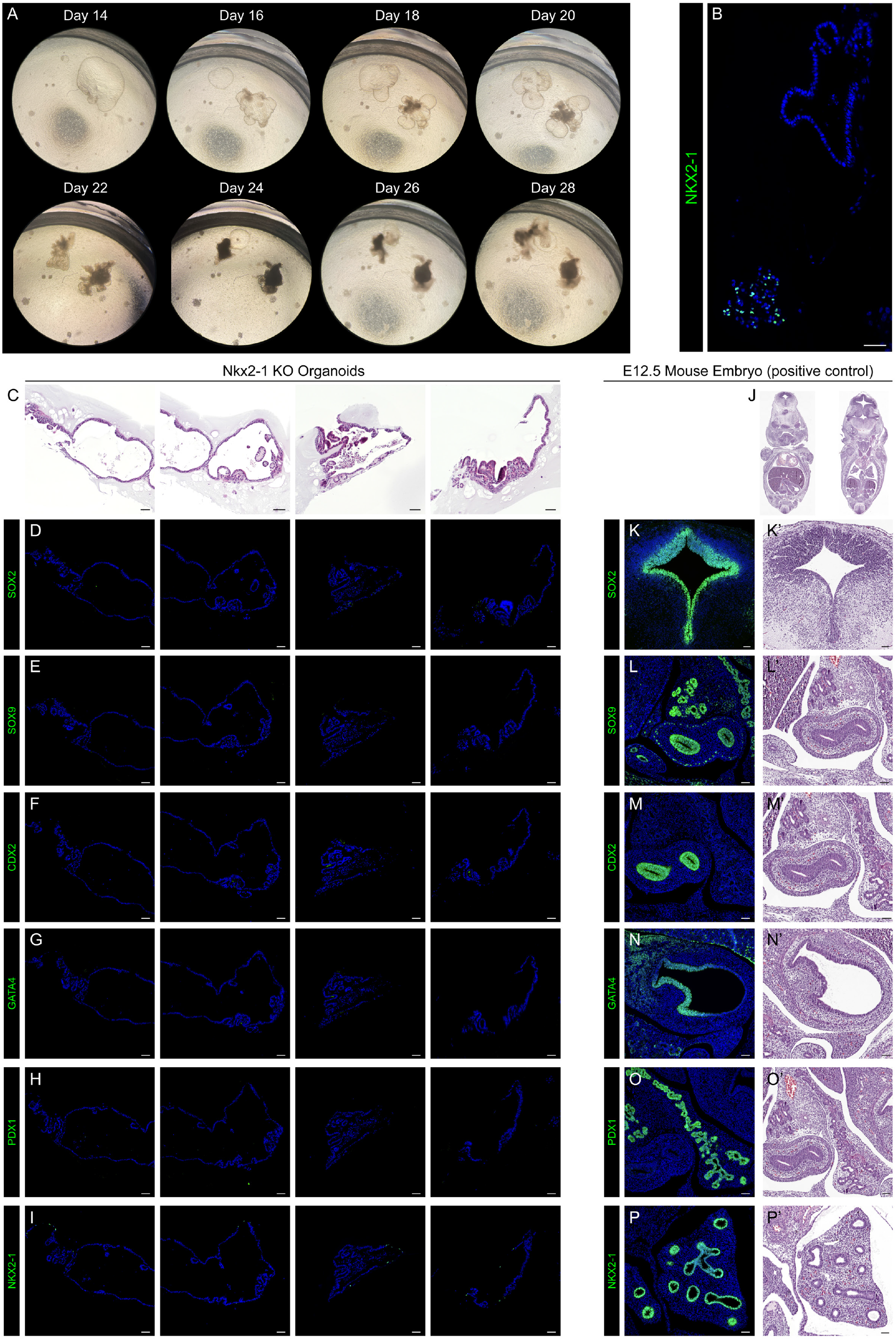
Nkx2-1 KO organoids lack substantial protein expression of canonical endoderm markers. (A) Time series of Nkx2-1 KO organoids in culture from day 12 to day 28. (B) Immunofluorescence of paraffin section from Nkx2-1 organoid Transwells showing non-recombined (Nkx2-1^+^) organoids adjacent to recombined (Nkx2-1^-^) organoids with atypical morphology. (C, J) H&E of paraffin sections of Nkx2-1 KO organoids (C) and E12.5 mouse embryos (J; K’-P’) used as positive controls for protein expression. Immunofluorescence of paraffin section from Nkx2-1 KO organoids (D-I) and E12.5 mouse embryos (positive controls; K-P) stained for canonical endodermal markers – Sox2 (D, K), Sox9 (E, L), Cdx2 (F, M), Gata4 (G, N), Pdx1 (H, O), and Nkx2-1 (I, P).

**Supplemental Figure 7.**
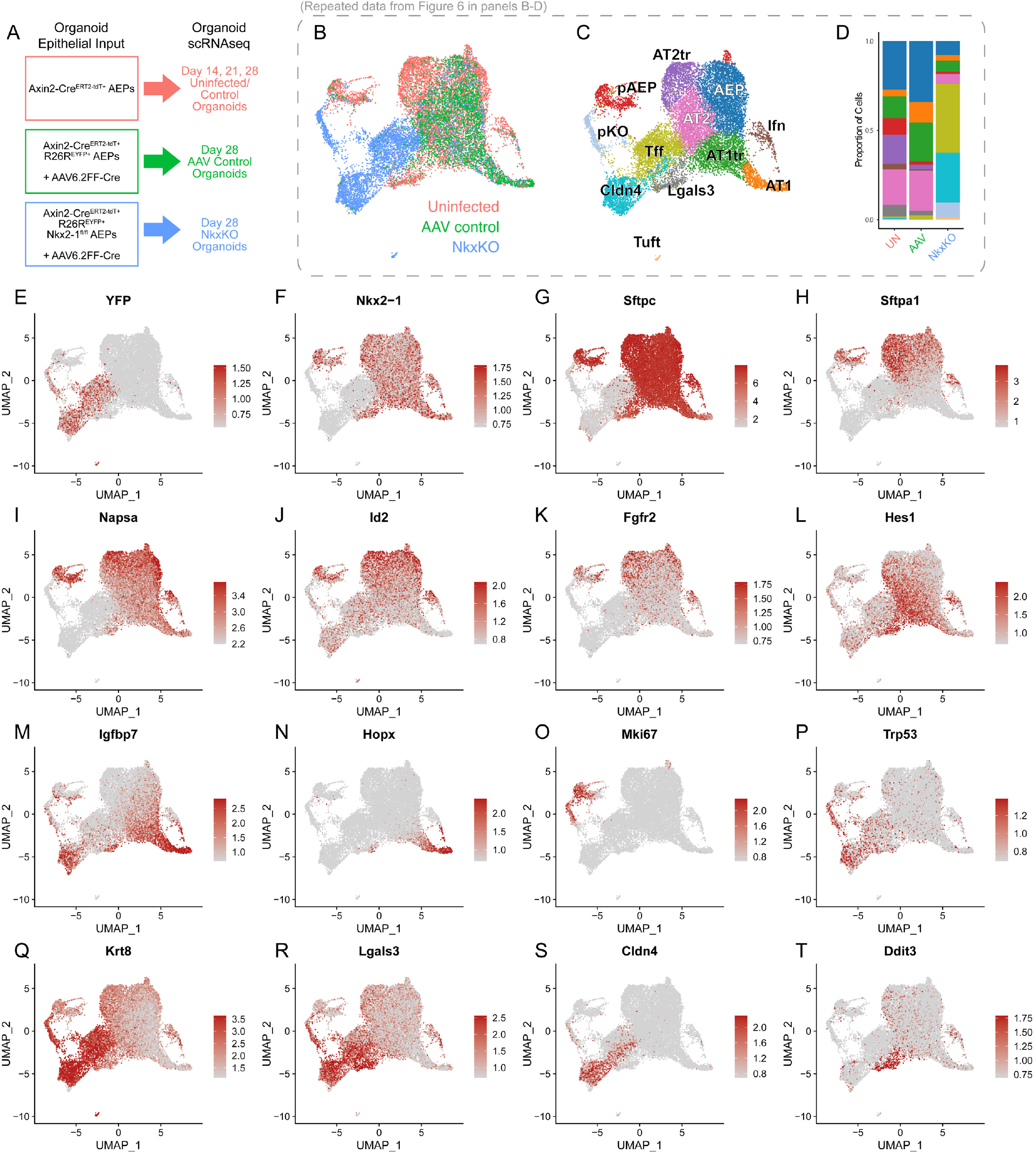
RNA Expression of Canonical Markers in Control (Uninfected), AAV Control, and Nkx2-1 KO AEP-derived Organoids. A) Overview of cell input used to generate AEP-O and subsequent timepoints for scRNAseq. (B-D) Repeated data from Figure 6K-M; Source (B), cell clusters (C), and proportion of cell type by experimental condition (D) for organoids outlined in (A). (E-T) RNA expression of common markers of alveolar epithelial cell identity.

## Notes

### Competing Interest Statement

The authors have declared no competing interest.

